# Neurodegeneration in human brain organoids infected with herpes simplex virus type 1

**DOI:** 10.1101/2021.03.05.434122

**Authors:** Agnieszka Rybak-Wolf, Emanuel Wyler, Ivano Legnini, Anna Loewa, Petar Glažar, Seung Joon Kim, Tancredi Massimo Pentimalli, Anna Oliveras Martinez, Benjamin Beyersdorf, Andrew Woehler, Markus Landthaler, Nikolaus Rajewsky

## Abstract

Herpes simplex virus type 1 (HSV-1) infection of the nervous system may lead to brain damage, including neurodegeneration. However, lack of suitable experimental models hinders understanding molecular mechanisms and cell-type-specific responses triggered by HSV-1. Here, we infected human brain organoids with HSV-1. Known features of HSV-1 infection such as alteration of neuronal electrophysiology and induction of antisense transcription were confirmed. Full-length mRNA-sequencing revealed aberrant 3’ end formation and poly(A)-tail lengthening. Single-cell RNA-seq and spatial transcriptomics uncovered changes in the cellular composition of the infected organoids caused by viral replication and dysregulation of molecular pathways in cell-type specific manner. Furthermore, hallmarks of early neurodegeneration were observed, namely extracellular matrix disruption, STMN2 and TARDBP/TDP43 downregulation, and upregulation of the AD-related non-coding RNA BC200/BCYRN1. These hallmarks were weaker/absent when infecting with a mutant HSV-1 control. Together, our data indicate that brain organoids serve as a powerful model to study mechanisms of HSV-1-driven neurodegeneration.

## Introduction

Herpes simplex virus type 1 (HSV-1) is a common human-specific pathogen affecting a large part of the population worldwide (1). After primary replication in epithelial cells, HSV-1 travels along axons innervating the affected regions to the trigeminal ganglia to establish a latent state of infection. The latent HSV-1 genome persists in episomal form within the nucleus, and HSV-1 DNA is chromatinized with heterochromatic histone marks. In this state, only a small subset of viral genes is transcribed, including the latency-associated transcripts (LATs) (2, 3). Investigations of post-mortem human tissue have provided evidence of LAT transcription/virus not only in trigeminal ganglia (4) but also in the brain at large (3). Latent viral infections can persist over years with very limited transcription. However, stimuli such as stress signals and weakened immunity can cause the reactivation of HSV-1 from sensory neurons at any time (3).

Two different forms of infection-related diseases in the brain have been observed. First, lytic infection, either by reactivation from latency or primary infection (5) in the central nervous system (CNS) can cause herpes simplex encephalitis (HSE), with an incidence of 1 in 10,000 infected individuals (6). HSV-1 infection in CNS is the most common cause of viral encephalitis (7). Second, HSV-1 is gaining increasing attention as a potentially causative agent in the pathogenesis of sporadic Alzheimer’s disease (AD) (8–12). Already decades ago, it was observed that HSE often occurred in the brain regions that are most frequently affected by AD (12). Furthermore, viral DNA was detected at high levels in amyloid plaques from sporadic AD post-mortem brain samples (13, 14). More recently, a population study reported that HSV-1 infection and seropositivity led to a significant risk of later development of AD (8,9,11,13,14). The current understanding of the mechanisms of HSV-1 infection, latency and reactivation in the CNS are based either on animal models or on *in vitro* differentiated neurons. These models cannot fully account for human-virus interaction specificity or cellular and functional diversity in neural tissues, respectively (15–18). Therefore, physiologically relevant human experimental models are needed to improve our understanding of the consequences of lytic and latent HSV-1 infection in the human CNS.

In recent years, human stem cell-derived brain organoids have emerged as models that capture several important aspects of human brain development and physiology (19–21). We hypothesized that brain organoids may offer an opportunity to molecularly study fundamental aspects of HSV-1 - brain tissue interaction. Indeed, recent data of acute and latent infections argue for the suitability of human induced pluripotent stem cell (hiPSC)-based neuronal models and brain organoids to study HSV-1 impact on neural cells (10,15,22). However, the CNS consists of many cell types, and it is unknown how efficiently the virus replicates in them, and whether these cell types respond differently to the infection. Interestingly, several features of AD, including amyloid plaque-like formation, gliosis and neuroinflammation were reported in 3D *in vitro* bioengineered neuronal culture (10). Nevertheless, the primary molecular mechanisms of HSV-1-driven neurodegeneration remain elusive but might be crucial for the potential identification of new drug targets.

In this study, we explored human cerebral organoids as a model system to capture cell-type specific molecular mechanisms employed by the virus and human brain cells to interact upon infection. Specifically, we used physiological and molecular bulk assays including full-length transcript sequencing (23) as well as single-cell RNA-sequencing and spatial transcriptomics to capture the functional and molecular consequences of HSV-1 infection on human brain cells in a physiologically relevant and complex cellular context and at single-cell resolution. Functionally, we were able to demonstrate with high-density multi-electrode measurements and, independently, real-time calcium imaging that HSV-1 infection massively damages neuronal activity in the organoids. Molecularly, we observed that HSV-1 heavily affected the cerebral organoid transcriptome at multiple levels, by activating antisense transcription, strongly modulating mRNA 3’ end formation and alternative polyadenylation, but also specifically dysregulating genes with synaptic, extracellular matrix and DNA-binding function. Furthermore, viral infection changed the cellular composition of organoids and triggered or repressed numerous cell-type specific molecular pathways, several of them being clearly related to neuroinflammation and early neurodegeneration. Specifically, we detected downregulation of neuronal growth associated protein stathmin-2 (STMN2) and TAR DNA-binding protein 43 (TDP43/*TARDBP*) and upregulation of the non-coding RNA BC200/*BCYRN1*, which was shown to be dysregulated in AD patients’ brains (24). Expression changes of these genes are associated with AD pathogenesis (24–26).

## Results

### Modeling HSV-1 infection using human cerebral organoids

We generated cerebral brain organoids from two different iPSC lines according to the optimized Lancaster et al. protocol (19,27,28) (Figure 1A). These organoids had predominantly dorsal forebrain region specification and contained multiple ventricle-like structures formed by SOX2/PAX6-positive neural progenitors surrounded by EOMES-positive intermediate progenitors and an early cortical-like structure formed by MAP2/TUJ1/LHX2-positive neurons (60 days old organoids, Figure 1A; 30 and 60 days old organoids, **Figure S1A**). At day 60, glial fibrillary acidic protein (GFAP) positive astrocytes started to emerge (**Figure S1A**). Also, the outer subventricular zone and neuronal layers expanded, which was reflected by expression of outer radial glia marker - *HOPX* and different neuronal markers such as TBR1/*TBR1* (deeper layer cortical neurons), CTIP2/*CITP2* and Reelin/*RELN* of Cajal-Retzius neurons (Figure 1B and **S1A, B**). These data indicate that cerebral organoids generate major cell types typical for early brain development. Furthermore, gene expression comparison between organoids derived from the two different iPSC lines showed an overall similar developmental signature, with the exception that iPSC line-1 additionally showed induction of the mesodermal lineage (*VCAN*, *LUM*), whereas iPSC line-2 showed more homogenous development towards dorsal forebrain (**Figure S1C**).

**Figure 1.**
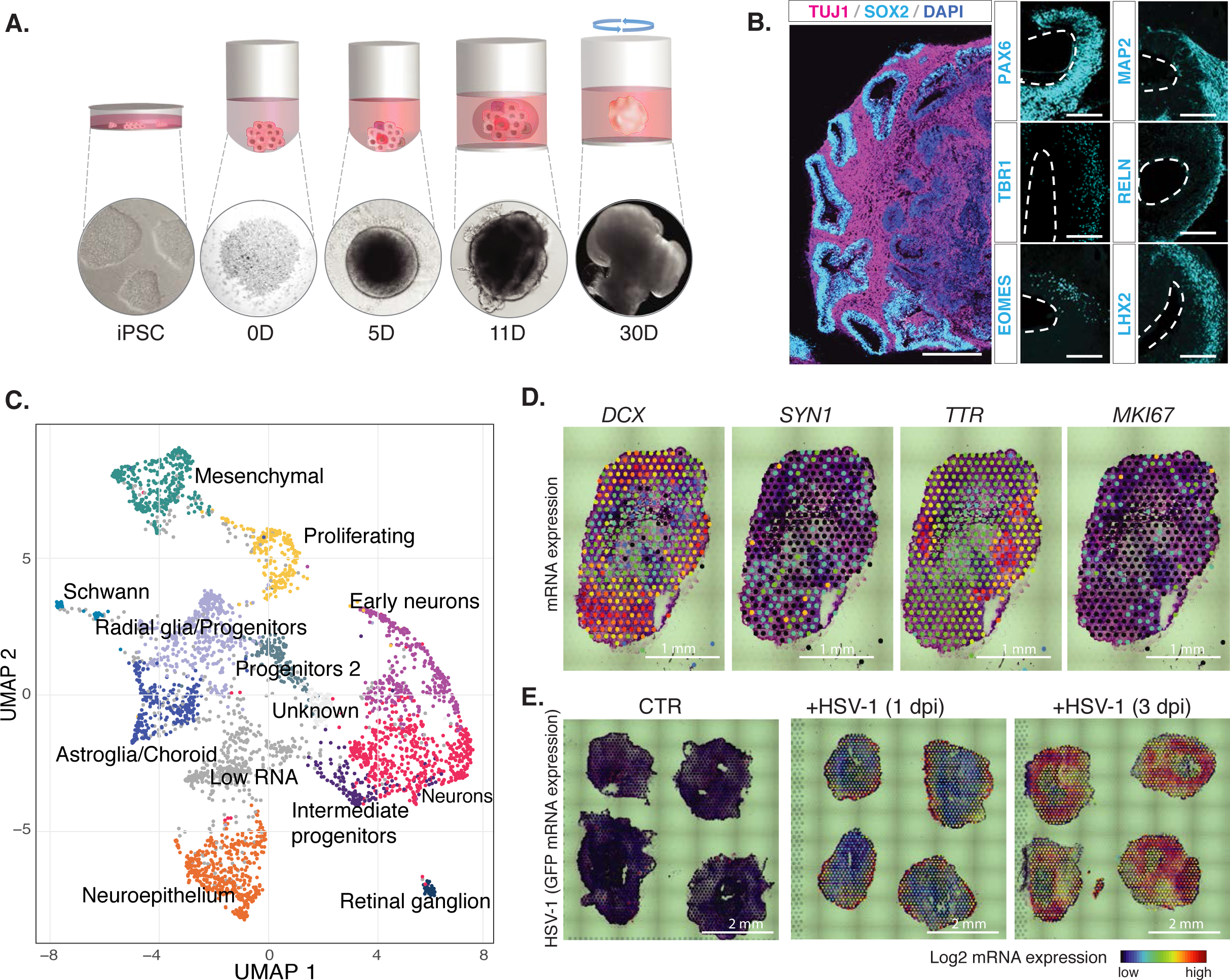
Human brain organoids as an experimental model to study mechanisms of HSV-1 lytic infection in complex brain-like tissue. **A**. Schematic overview of cerebral organoids generation using iPSCs. Lower panel: representative images of organoids at different developmental stages. **B.** Exemplary immunohistochemistry of 60 days old uninfected brain organoid showing location of neuronal progenitors (SOX2-cyan) and neurons (TUJ1-magenta). Scale bar - 500 µm Small panels display higher magnification of representative immunohistochemistry for progenitors (SOX2-cyan), intermediate progenitors (EOMES-cyan), neurons (MAP2, RELN, LHX2-cyan), deeper layer cortical neurons (TBR1-cyan) and nuclei (DAPI-blue). Scale bar - 100 µm. **C.** UMAP analysis of cells colored by cell type in uninfected 60 days old brain organoid. **D.** Representative images from the spatial transcriptomics analysis of 60 days old organoids for markers: *DCX* - neurons, *SYN1* – synapses, *TTR -* choroid plexus*, MKI67* - proliferating radial glia (RG). Colors indicate log2 mRNA expression levels. **E.** Spatial transcriptomics analysis of HSV-1 GFP expression in uninfected (CTR) as well as 1 and 3 days post infection (dpi). Colors indicate log2 GFP mRNA expression.

To assess cell type composition of developing organoids, we performed single-cell RNA-seq analyses of 30 and 60 days old organoids (Figure 1C and **S1D, E**). We combined highly reproducible single-cell transcriptomes from two biological replicates (60 days shown in **Figure S1D**) of each time point and identified six and fourteen transcriptionally distinct clusters in 30- and 60-day-old organoids, respectively (Figure 1C and **S1E**). Additionally, we performed differential expression analysis to select genes which defined each cluster (examples **Figure S1F**). In this way, we were able to identify multiple types of progenitors and neurons in 30 days old organoids and additionally astroglia in 60 days old organoids (Figure 1C and **S1E**).

Furthermore, we used the 10x Genomics Visium platform to define spatial mRNA expression in 60 days old organoids derived from both iPSC lines. We checked the distribution and colocalization of different cell-type specific markers using the Loupe Browser (10x Genomics). Neuron-specific transcripts such as the pan-neuronal marker *DCX* and synaptic marker *SYN1*, colocalized in several regions around the organoid periphery, while a marker of proliferating progenitors (*MKI67*) localized in regions compatible with ventricle-like structures. Also, *TTR*, a marker of choroid plexus, showed localized expression (Figure 1D for iPSC line-1 and **S1G** for iPSC line-2). These results correlate well with the immunohistochemistry of developing cerebral organoids (19).

Next, we infected 60 days old brain organoids with HSV-1-GFP and performed a variety of analyses shown in **Figure S1H** to investigate molecular, cellular and physiological changes. First, however, we assessed the infection efficiency by spatial transcriptomic analysis of GFP mRNA expression and by immunohistochemistry for GFP protein (Figure 1E and **S1I**). We detected viral RNA throughout the entire organoid, reaching even the deepest inner layers already at 3 dpi (Figure 1E), demonstrating the viral spreading from exterior to interior. On the protein level as measured via the GFP signal, we observed efficient spreading of the virus into the organoid outer layers at 3 dpi and expanding into the deeper inner layers at 6 dpi (**Figure S1H**). As expected, the necrotic core of the organoids did not show a signal for mRNA or protein GFP expression (Figure 1E **and S1I**).

In order to assess transcriptome changes of infected organoids, we performed a systematic analysis of bulk RNA expression, single-cell RNA-seq analysis and 10x Genomics Visium for all time points (3 and 6 dpi) and experimental conditions. Furthermore, we assessed the changes in alternative polyadenylation and poly(A) tail length by full-length mRNA sequencing and spontaneous neuronal activity by extracellular high-density multielectrode array (MEA) recordings and calcium imaging after HSV-1 infection. Immunostainings and real-time quantitative PCRs served as validation tools (**Figure S1H**).

### Molecular analyses of HSV-1 infected organoids reflect known facts about HSV-1 biology

To identify genes related to virus response, we profiled the transcriptomes of 60 days old uninfected, and infected (3 dpi and 6 dpi) organoids from two iPSC lines using bulk RNA-seq. Because a large fraction of sequencing reads from the infected samples aligned to the HSV-1 genome (up to 70%), we discarded these reads when analyzing cell types by clustering, since viral gene expression would otherwise dominate the analyses. In the principal component analysis, the first component reflected the genetic background and a slightly different developmental pattern of the two iPSC lines (**Figure S2A**). The second component indicated strong deregulation of gene expression as a consequence of HSV-1 infection. Differential gene expression analysis was designed to account for the differences between genetic backgrounds (principal component (PC)1), and measure the consistent effects of infection (PC2) on gene expression (see Methods).

This analysis of pooled samples from two iPSC lines showed that the infection led to systematically and statistically significant (p-value <0.05, log2 fold change cutoff 1) increased or decreased expression of 274 and 230 genes, respectively (**Figure S2B**). Downregulated genes were enriched for genes involved in synaptic function, extracellular matrix organization and development (**Figure S2C** and **D**), while upregulated genes were significantly enriched for DNA-binding transcription factor activity (adjusted p-value < 0.05) (**Figure S2E**). These gene expression changes were consistent between two iPSC lines, as shown in **Figure S2D** and **E**.

It has been previously reported that lytic HSV-1 neuronal infection causes synaptic dysfunction, associated with disassembly of key structural components of dendritic spines and upregulation of activity-regulated cytoskeleton-associated protein (ARC) mRNA and protein expression (29, 30). In our system, we observed massive downregulation of major pre- and postsynaptic components, including synaptic vesicle genes (*SV2A*, *SV2B*, *SYP, VAMP2*) and genes that function in calcium signaling (*SYT13*, *CAMK2A*, *CAMK2B*, *CALB2*) (Figure 2A). On the other hand, at the early stages of the infection, we could observe the previously reported upregulation of neuronal activity-related genes such as pentraxin *NPTX2, EGR2* or *ARC* (24). The expression changes for synapsin-1 (SYN1), postsynaptic density scaffolding protein 1 (HOMER1) and ARC were also shown for the proteins using immunostaining (Figure 2B) and western blot analysis (Figure 2C).

**Figure 2.**
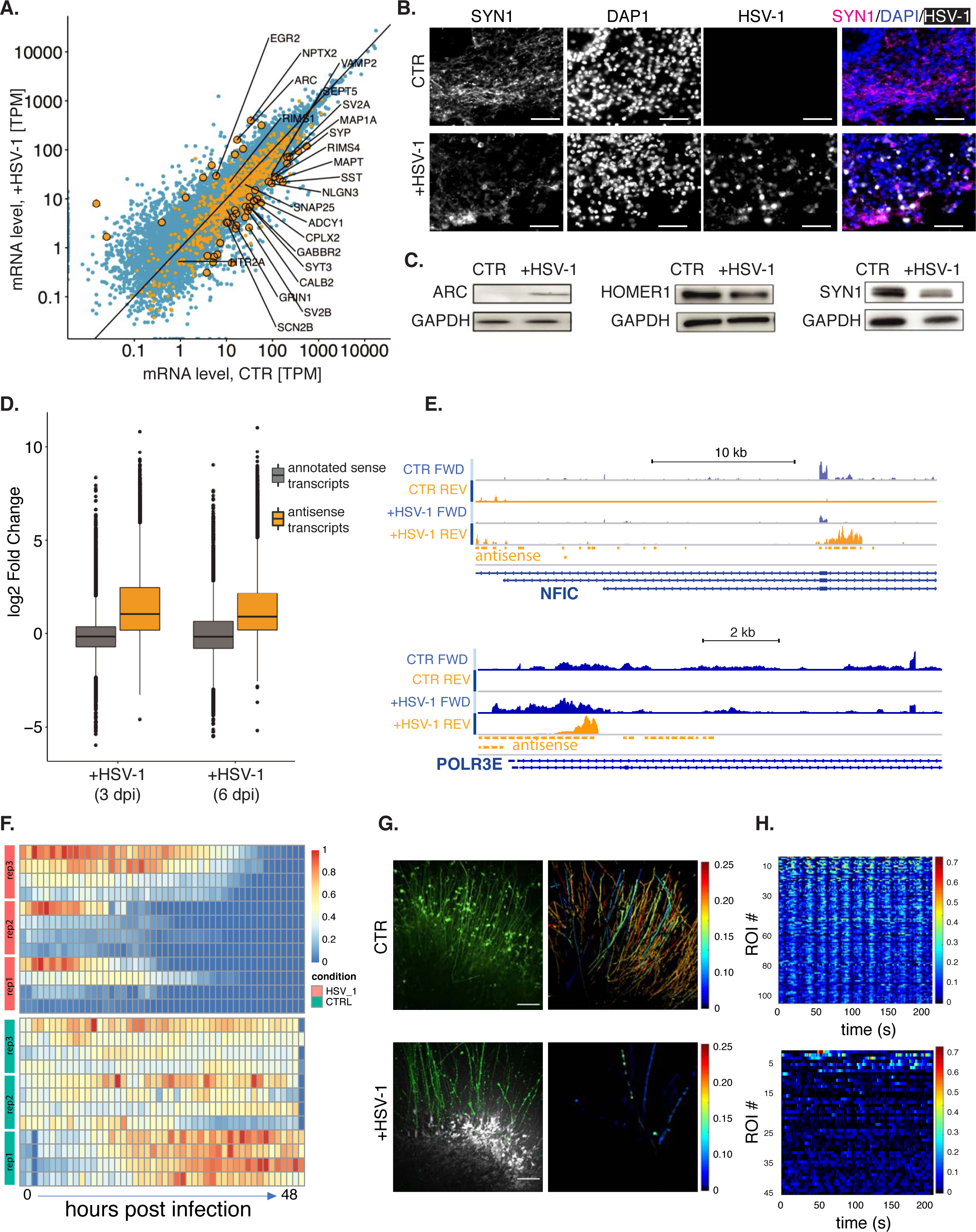
Infection of brain organoids captures known facts about HSV-1 infection: impaired synaptic activity and activation of antisense transcription. **A.** Gene expression in uninfected (CTR) and 3 days infected (+HSV-1) organoids. Yellow: transcripts related to chemical synaptic transmission. Large data points: statistically significantly de-regulated genes. Blue: other transcripts. **B.** Exemplary immunohistochemistry for synaptic (SYN1 – white, magenta) neuronal, nuclei (DAPI – white, blue) and HSV-1 (GFP - white) in uninfected and infected organoids. Scale bar - 100 µm. **C.** Representative western blots of synaptic proteins (ARC, HOMER1, SYN1) and a housekeeping protein (GAPDH) of uninfected (CTR) and infected (+HSV-1) organoids. **D.** Box plots showing log2 fold changes of sense annotated genes (grey) and antisense transcripts 3 and 6 days post infection (dpi). **E.** Examples of antisense transcripts: *NFIC* (antisense internal) and *POLR3Eas* (antisense divergent). Coverage profiles of total RNA-sequencing data from 60 days old cerebral organoids. Sense genes in *blue* (orange) oriented *left* to *right (right to left).* Transcript annotation in *dark blue*. **F.** Heatmap plots of the number of spikes captured by the most active electrodes of each replicate (n=3) in uninfected (CTR, green) and HSV-1-GFP infected (+HSV-1, red) organoid over 48 hours. **G.** Representative image of CalBryte590 AM (green) loaded, showing GFP virus expression (grey) at 48 hours post infection (hpi). Scale bar – 100 μm (left panel). Color-coded images illustrating spike frequency (right panel). **H.** Spike detection plot for all regions of interest (ROI) in uninfected (CTR – upper panel) and HSV-1-GFP infected (+HSV-1 – lower panel) organoids.

On the transcriptome level, we could recapitulate a widespread induction of antisense transcription, which has been previously reported in human primary fibroblasts (31). For the detection of antisense transcripts in 60 days old organoids, we defined all transcribed regions outside of annotated genes using the running sum algorithm (31) and filtered out potential transcriptional readthroughs. In total, we defined 5279 antisense transcripts. Detected antisense transcripts showed strong and statistically significant upregulation upon HSV-1 infection at both observed timepoints (Figure 2D). Selected examples of antisense transcripts are shown in Figure 2E.

### HSV-1 infection of brain organoids causes decreased synaptic firing rates and loss of coordinated neuronal activity

To validate that the observed changes in synaptic gene expression had functional consequences, we performed functional analyses using two independent approaches: high-density multielectrode array recordings and real-time calcium imaging (32–34). In the first approach, 60 days old organoids were placed on a 64-microelectrode array, which measures the voltage change from the large number of neurons present in the organoid tissue over time. Twenty hours after adding HSV-1 to the medium, the organoids showed a drastic decrease in the number of spikes compared to uninfected organoids, which maintained or even increased their spiking activity over time. No activity was detectable in the infected organoids after 40 h (Figure 2F and **S2F**). Brightfield microscopic images taken 48 h after infection showed that infected organoids also lacked outgrowing neurons, typically observed in uninfected organoids. (**Figure S2G**).

In the second approach to study the impact of HSV-1 on neuronal activity of infected organoids, we used calcium imaging (Figure 2G, H and **S2H, I**). To assess intracellular calcium dynamics of spontaneous action potentials in both uninfected and infected organoids, we performed CalBryte590 AM-based calcium imaging using spinning disk confocal microscopy. After plating the organoids, neurons (TUJ1-positive cells), showed active migration and neurite extension towards the glass dish bottom (**Figure S2H**). Calcium imaging analysis indicated a high abundance of spontaneously active neurons in all organoids (day 0). Next, we tracked changes in spontaneous activity after 24 and 48 hours post infection (hpi) with the HSV-1. Uninfected organoids reveal a high number of active neurites at 48 hpi present (Figure 2G and **Movie S1**), which show coordinated activity (Figure 2H). In contrast, in HSV-1 infected organoids, the robust initial spontaneous calcium activity decreased after 24 h and was almost completely abolished after 48 h in infected organoids (Figure 2G and H and **Movie S2**). After 48 h, the number of neurites was clearly reduced (**Figure S2H**) and the remaining neurites were silent or showed limited and uncoordinated activity (Figure 2I).

### HSV-1 infection causes global changes in poly(A) tail length and increased usage of distal polyadenylation sites

Previous studies reported that alternative polyadenylation may play an important role in antiviral response for viruses such as vesicular stomatitis virus (35) and that herpes infection can affect mRNA 3’ end formation by inducing premature polyadenylation (36) and readthrough transcription (37). Thus, to analyze the impact of herpes infection on alternative polyadenylation and, possibly, on poly(A) tails, we applied our recently developed full-length mRNA and poly(A) sequencing method (FLAM-seq, (23)) on three biological replicates of uninfected and HSV-1-infected 60-day-old organoids.

FLAM-seq allows to (a) exactly annotate mRNA 3’ ends and (b) measure the poly(A) tail length of mRNAs. Read counts for cellular and viral genes had high correlation among replicates (**Figure S3A).** All tail length measurements and 3’ end annotations are reported in **Tables S1** and **S2**. We observed that viral transcripts had poly(A) tails with a similar length distribution of cellular transcripts (**Figure S3B**), suggesting similar kinetics of tail synthesis and deadenylation. However, these distributions are mildly but significantly longer (median of 158 nt taking in to account the GFP transgene, 139 nt without, compared to a median of 122 nt in uninfected organoids, p-value = 2.2E-16, distributions shown in **Figure S3C**). Also, the median tail lengths of cellular transcripts increased significantly upon infection (from a median of 122 nt to 135 nt, p-value = 2.2E-16). While the overall effect of infection was to increase tail lengths of cellular mRNAs, we note that 116 mRNAs had a significantly shorter tail (threshold of 25 nt, p-value < 0.05, Figure 3A and **S3D**). Interestingly, genes with shorter tails were significantly enriched in functional categories directly linked to viral infection (“positive regulation of viral transcription” and “regulation of viral transcription”), while mRNAs with longer tails were not enriched in specific functional categories. Since mRNAs that are highly translated tend to have short poly(A) tails, we hypothesize that tail lengthening might be linked to reduced translation of cellular mRNAs, while tail shortening of transcripts that are important for viral replication might reflect higher engagement in translation. We conclude that HSV-1 infection causes a global deregulation of poly(A) tail lengths in cellular transcripts, possibly reflecting their different transcriptional and translational status.

**Figure 3.**
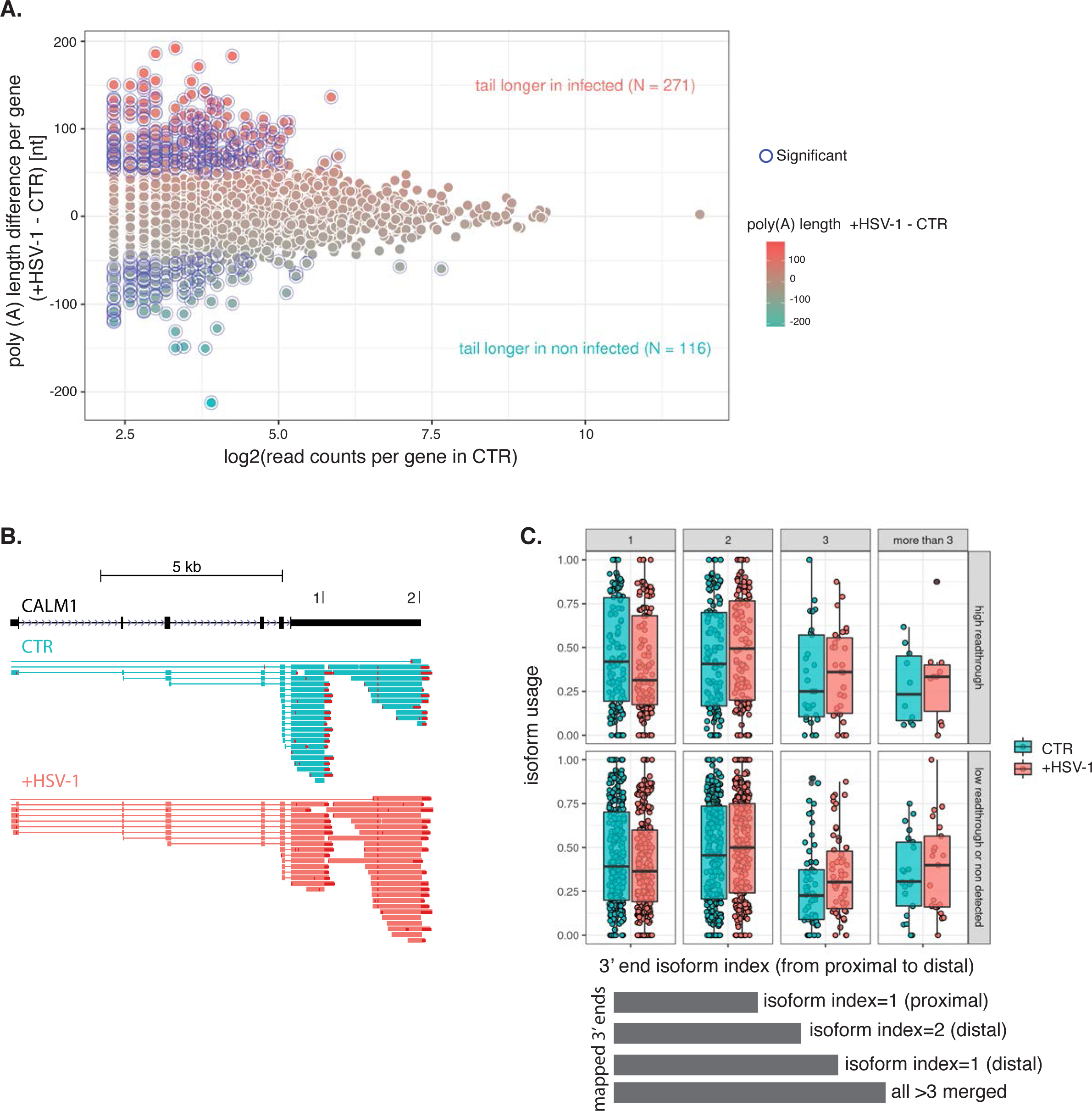
HSV-1 causes genome-wide lengthening of human poly(A) tails and abundant changes in polyadenylation site usage. **A.** Difference of median poly(A) tail length per gene in infected minus uninfected (in nucleotides), plotted against log2 expression level in uninfected (in read counts). Fill color represents the same as the y-axis to better visualize the changes in tail length. Blue circles mark genes with different median length of at least 25 nt and p-value < 0.05 (n = 1, three merged replicates, see methods). **B.** An example of shift in TTS usage, after HSV-1 infection for calmodulin 1 (*CALM1)*. The FLAM-seq reads from uninfected organoids are depicted in green, and from infected organoids in red. Poly(A) tails are appended to the 3’ end of reads and visualized as stretches of mismatches in red. Gencode annotation for the longest isoform is shown above in *black*. **C.** Isoform usage in infected (red) and uninfected (green), sorted by 3’ end isoform index, from proximal to distal. A scheme of isoform index annotation is provided below. The change in isoform usage between non-infected and infected was tested with Wilcoxon rank sum test (for proximal isoforms in CTR vs HSV-1, p-value = 0.0098). The difference in the change in proximal isoform usage between high readthrough genes and other genes was tested with Wilcoxon rank sum test (p-value = 0.2548).

To study possible changes in polyadenylation site usage, we annotated transcription termination sites (TTSs) supported by at least two reads and labeled 3’ end isoforms for each gene with an “isoform index”, equal to 1 for the most proximal isoform and increasing distally. For several mRNAs, we observed a striking difference in usage of distal TTS in infected organoids, as shown for calmodulin 1 (*CALM1*) (Figure 3B).

The decrease of the usage of the first, proximal isoform and an opposite increase in the usage of all downstream isoforms were global and significant in the infected organoids (p-value = 0.0098, Figure 3C and **S3E**), indicating a global shift in alternative polyadenylation upon HSV-1 infection. Since it has been observed that HSV-1 infection induces readthrough transcription in cellular genes, we asked whether the observed shift in TTS usage could be caused by failure in cleavage and polyadenylation, which could in turn increase the probability of downstream polyadenylation signal (PAS) usage. We therefore stratified the sequenced isoforms by published data (38) on HSV-1-induced readthrough transcription and observed that genes with high readthrough experience a slightly larger shift from proximal to all distal isoforms usage upon infection than genes with low readthrough, although this difference was not significant (p-value = 0.2548, Figure 3B and **S3E**). We therefore conclude that viral infection induces a global shift of 3’ end isoform usage from proximal to distal termination sites. Whether the observed shift is linked to transcriptional read-through or to alternative, unknown mechanisms warrants further investigation.

### Infection changes the cell-type composition of organoids

Using droplet-based single-cell RNA-sequencing (39), we performed single-cell transcriptome profiling of 60-day-old organoids at 3 and 6 dpi, capturing both host mRNA and viral transcripts. Since the RNA levels at 6 dpi were very low, likely due to extensive host cell shutoff (40), we focused on the 3 dpi time point in further analyses. An overview of the dataset (**Figure S4A**), with the number of characterized cells for each sample, distribution of unique molecular identifiers (UMIs), i.e., the number of individually detected mRNA molecules per cell, the number of detected genes and the mitochondrial mRNA counts are provided (**Figure S4B**). To normalize and scale the gene expression values to account for differences in sequencing depth as well as align cells with similar transcriptome profiles across conditions, we used the scTransform method (41).

By matching the transcriptomes of the individual cells to known marker genes, we defined 14 cell types in both uninfected and infected organoids, including progenitor cells, astroglia, intermediate progenitor cells, mesenchymal cells, terminally differentiated neurons, and low RNA/highly infected cells (Figure 4A, B and **S4C, D**). Within the infected organoids, the latter correspond to highly infected cells or cells with advanced HSV-1 infection stage (Figure 4A, B).

**Figure 4.**
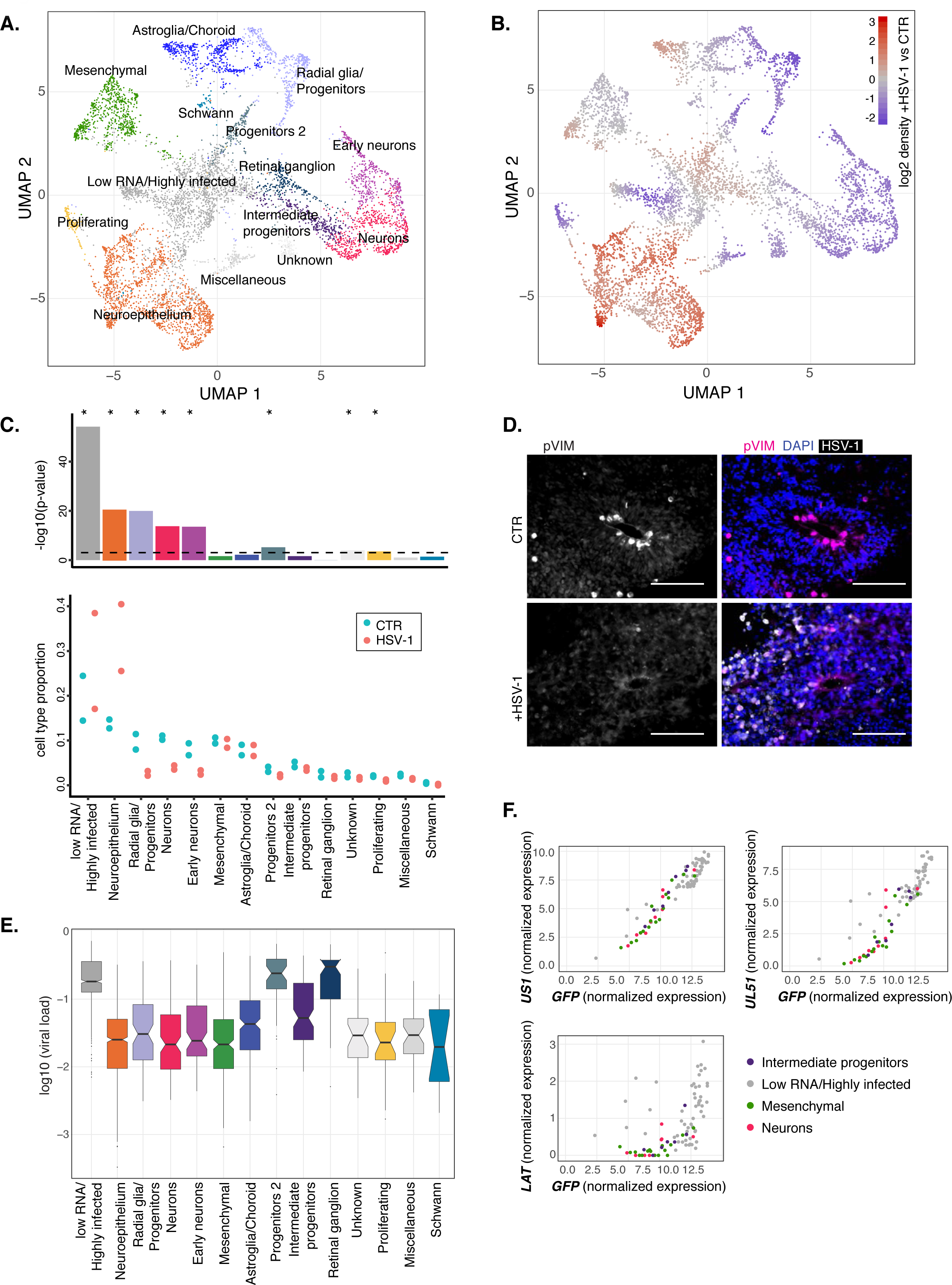
Cell-type specificity of virus infection susceptibility. **A.** and **B.** UMAP embedding of cells from uninfected organoids and 3 days post infection (dpi). In (**A**), cells are colored by their respective type. In (**B**), cells were colored by the relative density (infected vs. uninfected) of the area in which they are located, with red and blue indicating increased and decreased density upon infection, respectively. Example: cells in the bottom left are in an area with increased number of cells from infected organoids. **C.** Proportion of cell types in uninfected or infected organoids (lower panel), and p-values for testing differences (upper panel). Dashed line indicates p-value cutoff of 0.001. **D** Representative immunohistochemistry for mitotic marker (pVIM – white) and merged images for mitotic marker (pVIM - cyan), HSV-1 (GFP - white), nuclei (DAPI - blue) in uninfected (CTR) and 3 dpi infected (+HSV-1) organoids. Scale bar - 100 µm. **E.** Box plot showing log10 transformed percentages of viral RNA per cell (viral load). Lower and upper hinges correspond to the first and third quartiles. Whiskers extend to a maximum of 1.5 times the distance between the first and third quartile. Outliers beyond are marked by single dots. Notches extend 1.58 * interquartile range divided by the square root of number of cells. **F.** Cells corresponding from the indicated cell types were sorted by amount of viral RNA, grouped into bins of 20 cells, and normalized expression gene expression values calculated per bin. Plotted are the expression values of the GFP mRNA originating from the viral genome against *US1* (top left), *UL51* (top right), and *LAT* (bottom right).

Cell-type proportion changed significantly for several cell types, including proliferating cells, early neurons, neuroepithelial cells, neurons, and progenitor cells (p-value < 0.001, Figure 4C). For the majority of the cell types, a significant decrease in cell number was observed after infection, only neuroepithelial cells and low RNA/highly infected cells showed a significant increase in relative cell number after viral infection (Figure 4C). Notably, a prominent decrease in the number of mitotic cells in the ventricular zone and in the whole organoid were observed in 3 dpi organoids generated from both iPSC lines (Figure 4D for iPSC line-1 and **Figure S4E** for iPSC line-2). Overall, these findings demonstrate that HSV-1 infection resulted in a decrease in the number of proliferating progenitors as well as more differentiated cell types. On the other hand, the infection increased the number of low RNA/highly infected cells. Interestingly, these cells are not only characterized by high viral transcript load but also qualitatively by expression of different types of viral genes, suggesting a more advanced infection stage (**Figure S4G**).

### Cell-type specific levels of viral replication

Next, we asked to what extent the different cell types within the organoids would show distinct susceptibilities to infection. Although at 3 dpi, almost all cells contained viral RNA (Figure 1E**, S4H, I)**, the distribution of the viral load (percentage of viral transcripts/UMIs per cell) showed clear differences between cell types. Retinal ganglia, progenitors, and low RNA/highly infected cells had the highest amount of viral RNA, with a median of up to 10-fold more than other cell types (Figure 4E). A previous single-cell RNA-seq analysis of a HSV-1 infected cell line (fibroblasts) showed that some host cell genes are induced by the infection and that this induction correlates with accumulating viral RNA per cell (42). We counted the intronic reads (quantifying transcriptional induction) for three of these genes, namely *BCL2L11*, *RASD1* and *RRAD*. Among the cell types, which showed the strongest increase of these intron counts upon infection, were again the progenitor cells (**Figure S4G**). Thus, these data argue that not only the viral load is highest in this cell type but also the impact on host genes.

Given the observed cell-type specific viral RNA load, we asked if not only viral load but also known, temporally ordered viral gene expression programs would be cell-type specific. To investigate the course of the viral gene expression cascade in the different cell types, we ordered cells according to the amount of GFP mRNA, which serves as a proxy for viral replication, and grouped them into bins of 20 cells. For each bin, we calculated the relative expression values of the immediate early viral gene *US1*, and the late genes *UL51* and *LAT*. We then compared the values for intermediate progenitors, mesenchymal cells and neurons with the low RNA/highly infected cell group (Figure 4E). The kinetic class of the genes shown here is reflected by the shape of the curve, with *UL51* and particularly *LAT* starting to accumulate at higher amounts of GFP mRNA compared to *US1*, which steadily accumulates with increasing replication.

All genes, and particularly *LAT*, reached highest counts in the low RNA/highly infected cells, which suggests that the late stage of the viral gene expression cascade is only reached therein. However, the accumulation of the *US1* and *UL51* depended on the amounts of GFP, and thus the temporal development of the gene expression cascade, was largely overlapping in intermediate progenitors, mesenchymal cells and neurons.

Altogether, our data indicate that the different cell types show variable susceptibilities to viral infection.

### In advanced stages of HSV-1 infection, cells cannot be mapped to known cell types

Within infected organoids, we identified a large population of cells that clustered together with ‘low_RNA’ cells from control organoids. In uninfected organoids, we termed these cells ‘low_RNA’ by virtue of the low number of RNA molecules captured per cell (**Figure S4I**) and the absence of shared marker genes, which hindered their assignment to a defined cell type. Interestingly, in 3 dpi organoids, cells co-clustering with ‘low_RNA’ ones, were in addition characterized by high expression levels of viral genes (**Figure S4F**). We considered these cells to have lost a recognizable cell identity as a consequence of advanced viral infection.

To investigate the spatial distribution of ‘low_RNA’ cells in uninfected organoids and ‘highly_infected’ cells in HSV-1 infected organoids, we integrated the spatial transcriptomic and the scRNA-seq datasets (**Figure S4J-L**). Following cluster label transfer from single-cell RNA-sequencing, we observed that in uninfected organoids, spots with high probabilities for ‘low_RNA’ were mainly localized in the center of the section (**Figure S4K**, bottom row, left), indicating that ‘low_RNA’ cells were belonging to the organoid necrotic core. Conversely, already at 1 dpi higher scores for this type of cells were also found in the outer layers of the organoids (**Figure S4K**, bottom row, middle), where also high levels of virus-derived GFP mRNA was localized (**Figure S4K**, top row, middle). In contrast, spots with high probabilities for neurons clustered in few, discrete spots (**Figure S4K**, bottom row).

To corroborate the observation that spots in the spatial transcriptomics experiments bearing high levels of virus are abundant in highly infected cells, we plotted for each spot from the 1 dpi infected organoids the GFP expression level against the probability for low RNA/highly infected (**Figure S4L**). As expected, high probability values for this type were observed mostly at spots with high levels of virus-derived RNA (**Figure S4L**, left), whereas high neuron probabilities were only present in spots with moderate levels of GFP-mRNA (**Figure S4L**, right). This confirms that increasing amounts of virus-derived RNA coincide with distribution of highly infected cells.

### Viral infection induces and shuts down molecular pathways in host cells

To further characterize the cell-type specific transcriptional responses to HSV-1 infection, we performed a differential gene expression analysis between uninfected and 3 dpi organoids in each cell type (Methods, Figure 5A). Interestingly, the cell types with the highest amount of viral RNA (Figure 4E) did not show the strongest transcriptional response. Instead, numerous pathways were deregulated in epithelial and mesenchymal cells, which do not show high levels of viral RNA (Figure 4E).

**Figure 5.**
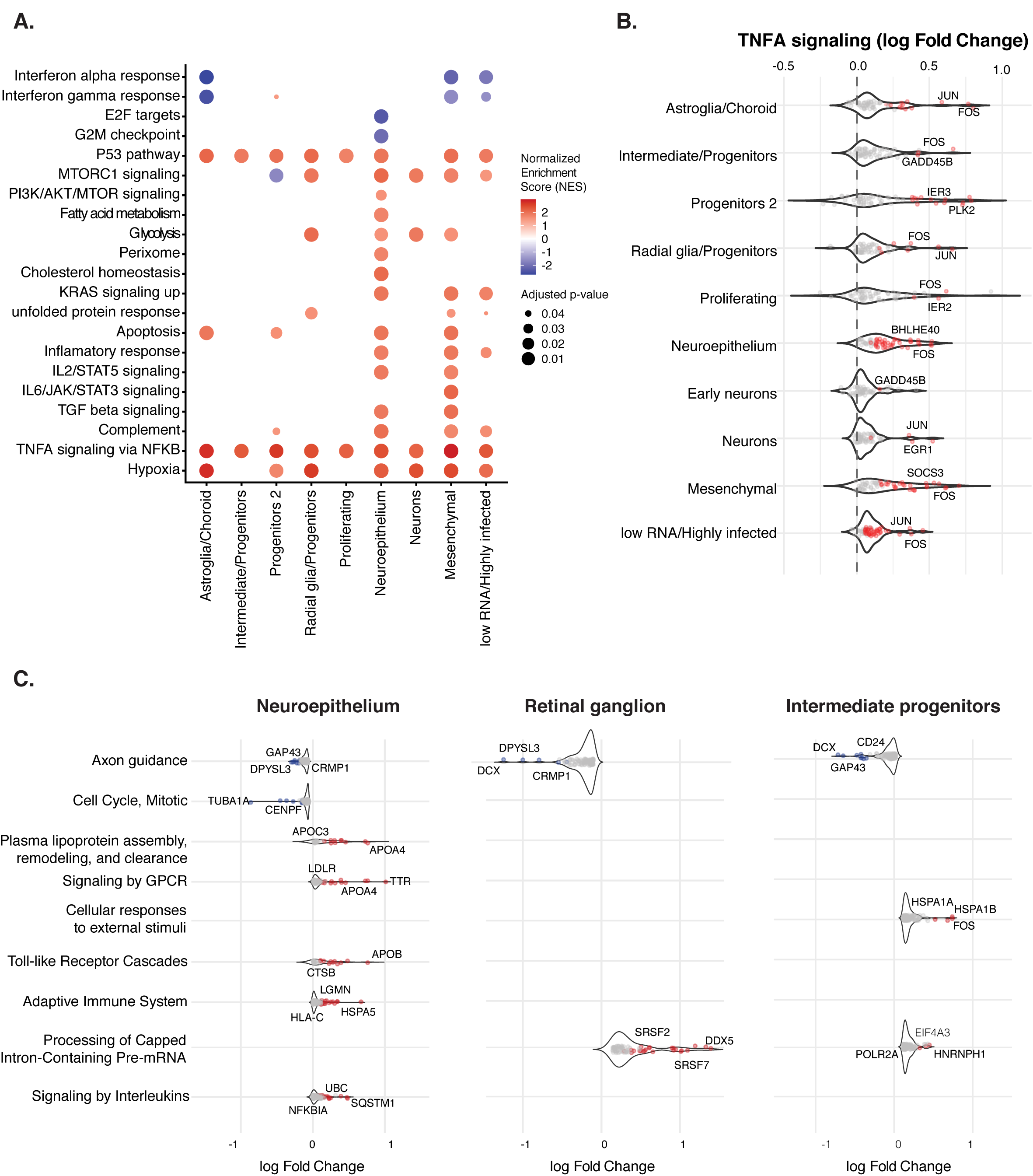
Viral infection induces and shuts down molecular pathways in host cells. **A.** Gene set enrichment analysis for each cell type, on differential expression between infected (3 days post infection (dpi)) and uninfected organoids. Color indicates normalized enrichment score (NES) values and dot size showing −log10 (FDR) from statistically significant pathways (FDR < 0.05). **B.** Violin plot showing log2 fold changes of TNF-α signaling for each cell type. Red dots: genes with positive log2 fold changes. **C.** Violin plot showing log2 fold changes for statistically significant genes per pathway for each cell type (neuroepithelium, retinal ganglion, intermediate progenitors). Red (blue) dots: genes with positive (negative) log2 fold changes.

We could identify pathways that were globally dysregulated in all cell types such as TNF-α signaling via NF-κB, hypoxia, and the p53 pathway (Figure 5A, B and **S5A**).

The enrichment of the p53 term was correlated with upregulation of genes such as *JUN*, *FOS*, *IER3*, *ATF3*, *BTG2*, *HEXIM1*, *NDRG1*, *SAT1* and *PLK2* (**Figure S5B**). A well-known tumor suppressor protein, p53 is activated by various stress signals. As viral infection evokes cellular stress, it is not surprising that infected cells activate the p53 pathway and also in our model we observed significant p53 activation in all cell types but, interestingly, not in neurons (**Figure S5B**).

Furthermore, we observed upregulated metabolism-associated genes. For example, the mammalian target of rapamycin complex 1 (*mTORC1*) pathway, which was upregulated in progenitors, neurons, neuroepithelium and mesenchymal cells but downregulated in the progenitors 2 cluster. HSV-1, produces capped mRNAs which engage cellular ribosomes. In order to translate its own mRNAs stimulates cap-dependent translation by activating mTORC1 to inhibit the translational repressor 4E-binding protein 1 (*4E-BP1*) (43, 44). Activation of *mTORC1* could therefore reflect an increase in the translation potential.

Additionally, neuroepithelial cells showed upregulation of lipoprotein metabolism and signaling by G-protein coupled receptor (GPCR) pathways, but downregulation of cell-cycle and axon guidance related pathways (Figure 5C). On the other hand, intermediate progenitor and retinal ganglion cells showed downregulation of axon guidance and upregulation of pre-mRNA processing related pathways (Figure 5C). Interestingly, these cell types contained a large amount of viral RNA, but their number was not affected upon infection (Figure 4C, F and **S4K)**. These results indicate that HSV-1 replication causes deregulation of different cellular pathways globally but also in a cell-type specific manner. Furthermore, our data suggest that individual cell types cope differently with viral replication and high viral RNA content is not always a prerequisite for the strongest transcriptional response.

### HSV-1 infection induces hallmarks of neurodegeneration

We investigated whether HSV-1 replication and inflammatory responses in infected organoids would lead to molecular changes associated with neurodegeneration. First, we assessed cell death by immunostaining with cleaved caspase-3 (CC3) antibody (Methods). We did not observe an increase in CC3 expression upon HSV-1 infection (**Figure S6A**). This is consistent with published reports from mouse models showing that herpes simplex viruses protect cells against apoptosis, via NF-κB activation and Casp3-dependent pathway (45). Next, we investigated early events which could lead to neuronal damage and synaptic dysfunction, preceding late pathological changes.

One of the most profound phenotypes observed after HSV-1 infection was disruption of the tissue integrity in infected organoids. Therefore, we analyzed the expression of extracellular matrix (ECM) components (Figure 6A**, S6B**). We observed a strong (more than four-fold) reduction of decorin (*DCN*), biglycan (*BGN*), versican (*VCAN*) and aggrecan (*ACAN*), and a mild effect (log2 fold change > −2) on hyaluronan production (*HAS1* and *HAS2*) in RNA-seq data after lytic organoid infection. We confirmed this finding by RT-qPCR analysis, using independent biological replicates for two different iPSC lines for collagen type 6A1 (*COL6A1*), desmin (*DES*) and *BGN* (Figure 6B). Furthermore, we validated a significant downregulation of collagen type 3A1 (COL3A1) by western blotting analysis (Figure 6C). On the other hand, we found heparan sulphate modifying enzymes such as 3-O-sulfotransferase 2 and 6 (*HS3ST2/HS3ST6*) among upregulated ECM transcripts (Figure 6A).

**Figure 6.**
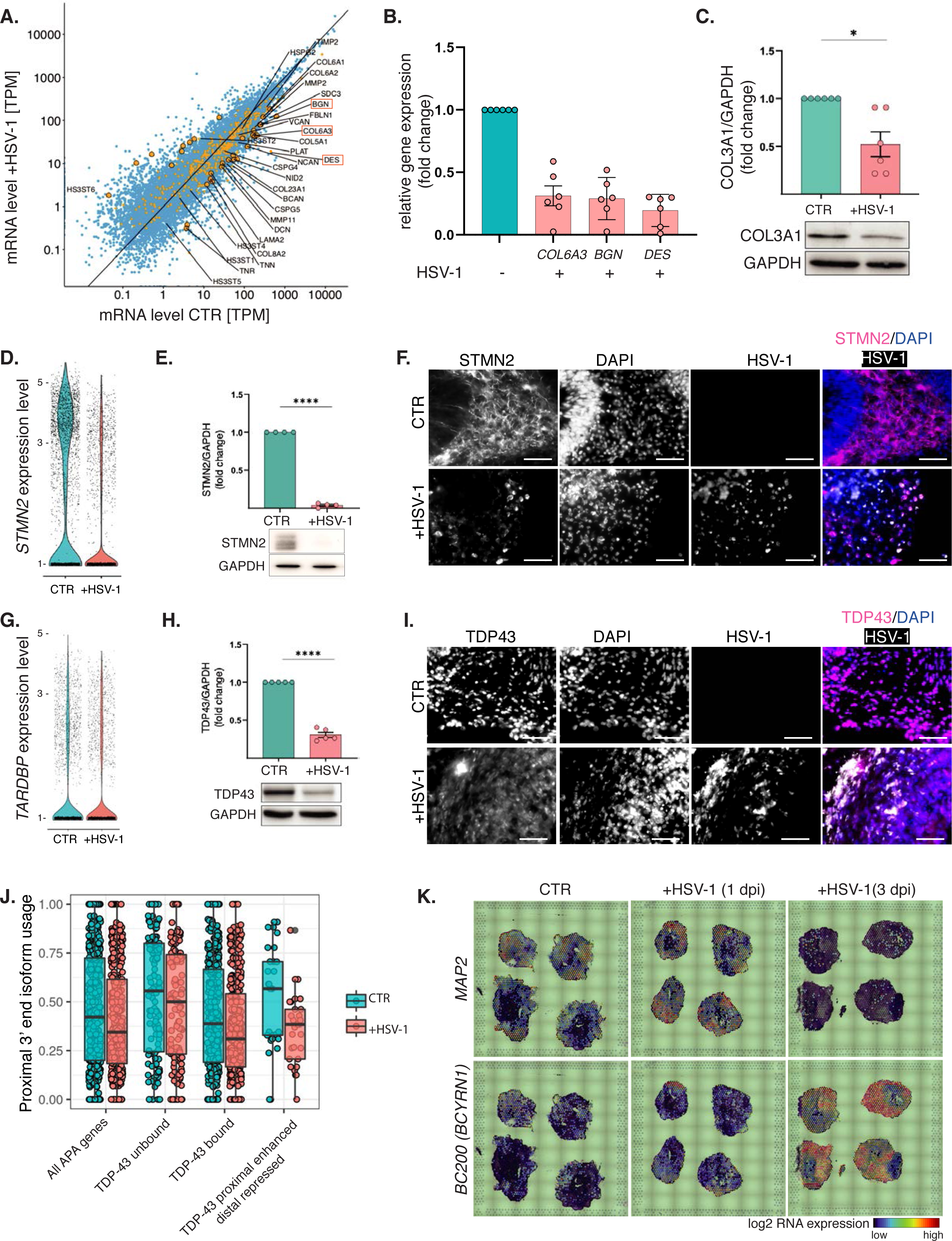
Hallmarks of neurodegeneration caused by HSV-1. **A.** Gene expression (transcripts per million - TPM) changes in uninfected (CTR) versus infected (+HSV-1) organoids, transcripts related to extracellular matrix (GO:0031012) in yellow, significantly deregulated ones show as large points, red boxes depict genes validated by qPCR. Blue data points indicate other transcripts. **B.** Quantitative real-time PCR analysis of *COL6A3, BGN* and *DES* transcript expression in organoids derived from two iPSC lines, 3 days post infection (3 dpi). Values are given as SEM, n = 6 biological replicates from two iPSC lines. Statistical analysis was performed by Wilcoxon rank sum test. *indicates statistically significant differences between uninfected and infected organoids (*p-value ≤ 0.05). **C.** Representative western blots and relative protein expression semi-quantified by densitometry analysis of collagen type 3A1 (COL3A1) of uninfected (CTR) and 3 dpi infected (+HSV-1) organoids, normalized to GAPDH expression. **D.** Single-cell mRNA expression of stathmin-2 (*STMN2*). Each dot represents a single cell. Statistically significant gene expression is observed only if a violin-shaped fitting area can be calculated. **E.** Representative western blots and relative protein expression semi-quantified by densitometry analysis of STMN2 of uninfected (CTR) and 3 dpi infected (+HSV-1) organoids, normalized to GAPDH expression. **F.** Exemplary immunohistochemistry for neurons (STMN2 – white/cyan), nuclei (DAPI - white/blue) and HSV-1 (GFP - white) in uninfected (CTR) and 3 dpi infected organoids (+HSV-1). Scale bar - 100 µm. **G.** Single-cell expression of TAR DNA-binding protein 43 (*TARDBP*). Each dot represents a single cell. **H.** Representative western blots and relative protein expression semi-quantified by densitometry analysis of TDP43 of uninfected (CTR) and 3 dpi infected (+HSV-1) organoids, normalized to GAPDH expression. **I.** Exemplary immunohistochemistry for nuclear protein (TDP43-red), nuclei (DAPI-blue) and HSV-1 (GFP-white) in uninfected (CTR) and 3 dpi infected organoids (+HSV-1). Scale bar - 100 µm. Values are given as mean ± SEM. n = 5-6. Statistical analysis was performed by unpaired student’s t-test for direct comparisons of two independent groups. *indicates statistically significant differences between uninfected and infected organoids (*p-value ≤ 0.05; ****p-value ≤ 0.0001) (two independent experiments). **J**. Proximal isoform usage (y axis) for genes with alternative polyadenylation. Of those, genes with proximal isoform not bound by TDP43 (TDP43 unbound), genes with proximal isoform bound by TDP43 (TDP43 bound), and genes where usage of proximal isoform was reported to be enhanced by TDP43 or distal isoform to be repressed by TDP43 in (50) (“Proximal favored by TDP-43”). Green dots and boxes represent non-infected samples, red infected samples. The different usage of proximal isoforms in uninfected organoid samples between TDP43-bound and unbound isoforms was tested with Wilcoxon rank sum test, p-value = 0.0012. When comparing the difference in proximal isoform usage of infected and non-infected samples between all the gene categories in x, only the last category differed significantly from the set of all genes with alternative polyadenylation (Wilcoxon rank sum test p-value = 0.0013). **K.** Representative images from the spatial transcriptomics analysis of 60 days old organoids for dendritic transcripts noncoding *BC200/BCYRN1* and *MAP2*. The colors indicate log2 expression.

Our analysis indicates that HSV-1 infection strongly affects expression of ECM components, which could trigger degeneration processes and could be associated with observed profound changes in organoid morphology and tissue integrity (**Figure S6C**).

The second most prominent phenotype accompanying destruction of the organoid structure was strong downregulation of stathmin-2 (*STMN2*) transcript in both bulk and single-cell RNA-seq data (Figure 6D). Consistently, STMN2 expression was significantly downregulated in HSV-1 infected organoids generated from both iPSC lines (Figure 6E, F and **S6D, E**). *STMN2* encodes a microtubule regulator, required for normal axonal outgrowth and regeneration. STMN2 downregulation has been associated with neurodegenerative diseases (46). It has been reported that STMN2 levels decrease following TAR DNA-binding protein 43 (*TARDBP/* TDP43) knockdown or relocalization from the nucleus in human motor neurons (47) and a direct link between TDP43 loss in patients, pathology, and STMN2 down-regulation (25). Therefore, we compared *TARDBP* mRNA expression in bulk and single-cell RNA-seq data, but did not observe significant downregulation (Figure 6G). Since *TARDBP* is known to be regulated post-transcriptionally, we analyzed TDP43 localization and abundance in uninfected and infected organoids. We observed clear nuclear localization of TDP43 in uninfected organoids and, upon infection, relocalization to the cytoplasm upon HSV-1 infection (Figure 6I for iPSC line-1 and **S6F** for iPSC line-2). Also, full-length protein level was significantly decreased (Figure 6H and **S6G**). Moreover, a known TARDP truncated 25 kDa neurotoxic C-terminal isoform (26) could be observed only in infected organoids (**Figure S6G**).

Besides splicing regulation, *TARDBP*/TDP43 is also involved in mRNA 3’ end formation (48, 49). Therefore, we tested whether its deregulation could play a role in the observed changes in alternative polyadenylation (as shown in Figure 3). We analyzed the presence of published *TARDBP*/TDP43 binding sites (46,47,50) at the 3’ ends of mRNAs that we had annotated from our full-length transcript sequencing data (Methods). It has been previously shown that *TARDBP*/TDP43 binding close to polyadenylation sites often, but not always, represses their usage (50). Consistently, we observed a significant decrease in the usage of proximal PAS in isoforms which were bound by *TARDBP*/TDP43 compared to isoforms which were not (p-value = 0.0012, Figure 6J). Given that viral infection affects the usage of proximal 3’ ends, and at the same time it affects *TARDBP*/TDP43 expression and localization, we would expect a possible attenuation in the downregulation of proximal isoforms. The difference in the isoform usage change upon infection between *TARDBP*/TDP43 bound and unbound 3’ isoforms was mild and non-significant (Figure 6J). On the other hand, for those genes that are known to be regulated by *TARDBP*/TDP43 in the opposite way (enhancing usage of the proximal PAS or repressing that of distal ones (50)) we observed that the viral infection induced a strong shift in 3’ isoform usage, significantly larger than the rest of the genes (p-value = 0.0013, Figure 6J). This is what would be expected knowing that *TARDBP*/TDP43 favors proximal isoform usage in these genes.

Furthermore, in our data we observed a dysregulation of non-coding RNA BC200/*BCYRN1*, which localizes to dendrites where it contributes to the maintenance of long-term synaptic plasticity (24). Although the majority of dendritic transcripts such as *MAP2* show downregulation upon HSV-1 infection, *BC200* was strongly upregulated already at 1 dpi, and its expression correlated with expression of HSV-1 viral genes (Figure 6K and **S4I**). We also confirmed this finding in the bulk RNA-seq data (**Figure S6H).**

Together, these results suggest that there are several molecular changes in HSV-1 infected organoids which could be linked to neurodegeneration and could contribute to later pathological changes.

### The Herpes simplex ICP27 mutant has weaker neurodegenerative potential

To determine whether the observed changes related to neurodegeneration are directly dependent on HSV-1 replication, we compared wild-type (WT) HSV-1 with a ICP27 deletion mutant (ICP27 MUT). ICP27 is an essential protein for the expression of late genes as well as the synthesis of viral DNA (46). Because the ICP27 mutant is known to be less efficiently replicating in human cells (51, 52), we compared organoids infected for two days with WT virus to these infected for four days with the mutant strain. This experimental setup allowed us to keep a similar expression level of viral proteins between conditions (Figure 7A). Because *STMN2* level was strongly affected by the WT virus, we performed western blot and immunostaining analysis of STMN2 expression in both conditions, to determine impact of the mutant virus on its expression. We observed that STMN2 was less depleted in ICP27 MUT when compared to WT and its protein expression anticorrelated with viral protein (ICP0) expression (Figure 7B, C). Also, the organoid integrity and proliferating progenitors were less affected by the mutant strain (**Figure S7A**). This suggests that the observed neurodegeneration related changes are directly related to viral replication and could be decreased (or perhaps even treated) by lowering the efficiency of viral replication as shown for HSV-1 ICP27 mutant.

**Figure 7.**
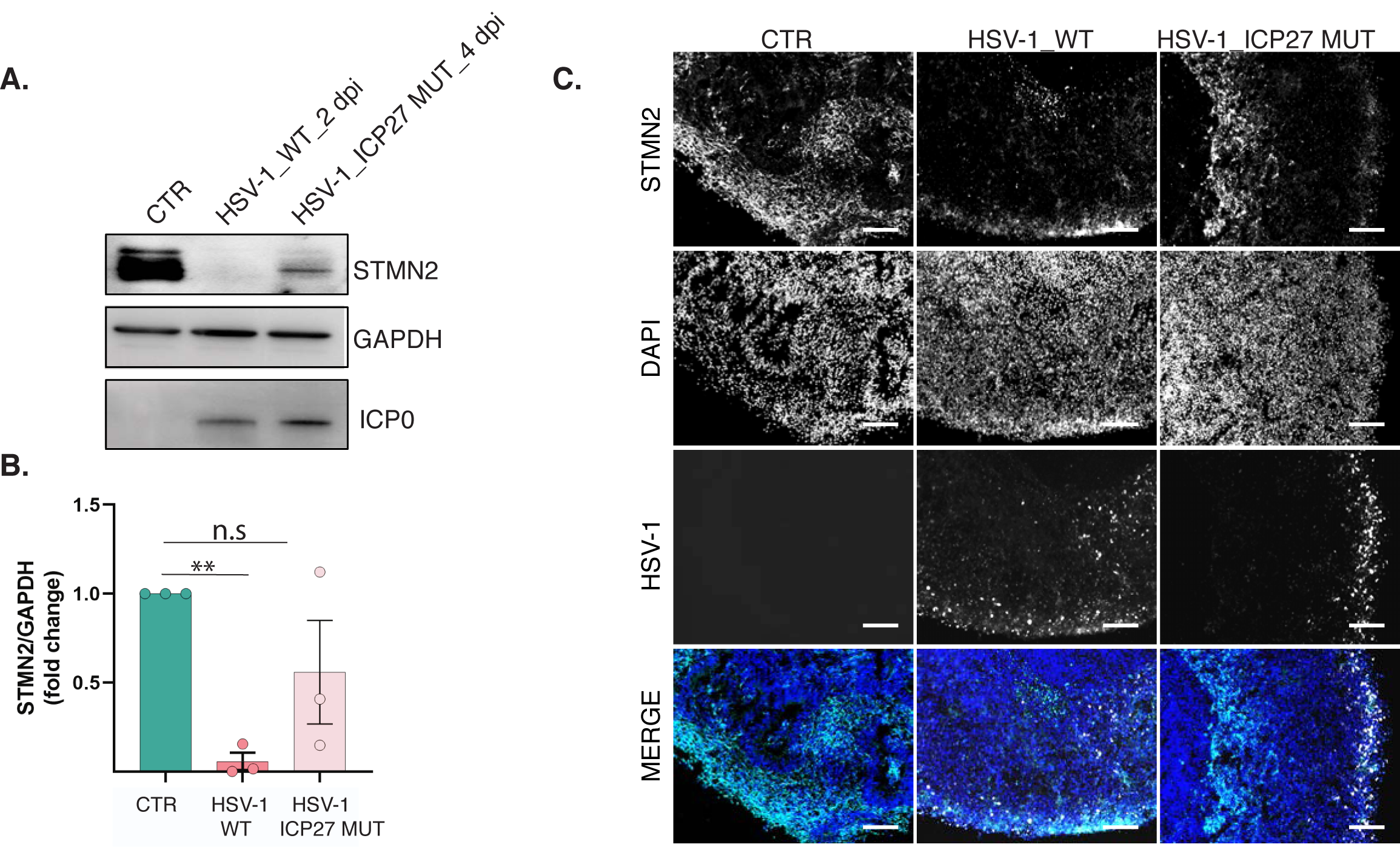
Herpes simplex ICP27 mutant exhibit weaker neurodegenerative potential. **A.** Representative western blots of stathmin-2 (STMN2), housekeeping protein (GAPDH) and viral ICP0 protein of uninfected (CTR) and organoids infected using HSV-1 wild-type (WT) virus (2 dpi), and ICP27 mutant (MUT) virus (4 dpi). **B.** Relative protein expression semi-quantified by densitometry analysis of STMN2 of uninfected (CTR), HSV-1 WT and HSV-1 ICP27MUT infected organoids, normalized to GAPDH expression. Values are given as mean ± SEM. n = 3. Statistical analysis was performed by unpaired student’s t-test for direct comparisons of two independent groups. *indicates statistically significant differences between uninfected and infected organoids (**p-value ≤ 0.01). **C.** Exemplary immunohistochemistry for STMN2 (red), DAPI (blue) and viral ICP0 (white) in the uninfected (CTR) and organoids infected with HSV-1 WT virus (2 dpi) and ICP27 MUT virus (4 dpi). Scale bar – 100 µm.

## Discussion

Although the course of HSV-1 infection has been studied, the molecular mechanisms of virus-driven brain inflammation and neurodegeneration are elusive. This is arguably due to the absence of human model systems that are experimentally accessible and that capture the complexity of different cell types linked together in neural tissues. In this study, we demonstrate that human brain organoids, combined with a broad range of RNA, protein, and electrophysiological analyses tools, can fill this gap. Specifically, we show **(a)** how HSV-1 “invades” neural tissue space, **(b)** how it differentially replicates in different cell types, **(c)** how it affects synapse functionality (impairing both spike rates but also spike coordination), **(d)** that it triggers the up-/down-regulation of specific human molecular pathways in a cell-type specific way, **(e)** that it targets certain molecular mechanisms such as adenylation site choice, poly(A) tail length, and antisense transcription to manipulate host-cell response, and **(f)** that it likely triggers the molecular onset of neurodegeneration (as we see the up-regulation/dysregulation of key markers of Alzheimer’s disease in infected organoids). We therefore believe that our results contribute to a much more detailed molecular and functional understanding of cell-type specific consequences of HSV-1 infection and establish human brain organoids as a powerful model system to advance this approach in the future. Of course, the brain organoids used in this study capture only a small part of the true complexity of human brains as they represent only fetal brains and lack immune cells and blood vessels. However, we did observe dysregulation of the interferon pathway (down-regulated in astroglia and mesenchymal cells) and the NF-*κ*B pathway (up-regulated in most cell types and thus blocking apoptosis, favoring HSV-1 replication). Both pathways are crucial for the function of the innate immune system, thus indicating that our organoid-infection model may be used to study how HSV-1 escapes the innate immune response in a neuronal tissue. In any case, the organoid field makes rapid progress and we believe that future organoid models will capture more of human brain biology (such as microglia) and thus, arguably, increasingly detailed and medically relevant aspects of HSV-1 infection. We believe that our study may serve as a reference point for this future research. Since organoids are genetically easily accessible and allow, as we demonstrate, to quantify cell-type specific effects of infection, perturbation studies may reveal new potential cell-type specific drug targets (or combinations thereof) to impact viral replication, or host-cell response, or possibly targets that modulate the onset of neurodegeneration. Moreover, organoids could serve as a model to screen for compounds, drugs, or modern vaccines, possibly in preclinical settings, that can prevent infection or treat the functional consequences of infection.

We now discuss one of the most striking molecular mechanisms that we found to be involved in modulating viral-host-cell interactions: alteration of poly(A) tails of cellular messages and a striking switch in their usage of polyadenylation sites, as revealed by our full-length mRNA and poly(A) tail sequencing. We note that we are not aware of any other animal system in which such a strong switch of 3’ UTR isoform usage has been reported previously, and believe therefore that this effect is particularly important for understanding the impact of HSV-1 infection on human cells. These tail alterations and 3’ UTR usage switches are highly likely to affect the expression, the localization and even the function of mRNAs and their resulting protein products. This is particularly obvious for the 3’ UTR usage switches as 3’ UTRs are key targets of (hundereds of) miRNAs and (thousands of) RNA binding proteins that alter localization, expression, turnover, and translation of the respective mRNA (53, 54). Regarding poly-A-tails, the de-adenylation of mRNAs is coupled with translation (55) and it is known that more actively translated and more stable mRNAs tend to have shorter poly(A) tails at the steady state (56). We observed a general increase in tail length of host mRNAs, which could reflect their reduced translation and/or different transcription and turnover kinetics. The observed shift in poly(A) length seemed to be rather global and did not affect any specific gene categories. On the other hand, fewer transcripts had reduced tail length upon infection. These were enriched for genes involved in promoting HSV-1 infection such as CDK9 (57) and could represent a subset of mRNAs crucial for early viral replication. Besides observing changes in poly(A) tail length of mRNAs, we report vast changes in alternative polyadenylation. It was recently shown that the viral protein ICP27 affects 3’ end formation of host mRNAs by binding the cleavage and polyadenylation complex and suppressing its activity on termination sites (58). The consequent transcriptional readthrough might lead to increased usage of distal polyadenylation sites, which we consistently observed upon infection. Interestingly, an opposite effect - a global shortening of 3’UTRs - was observed after vesicular stomatitis virus (VSV) infection in microphages (35), which suggest that effects on 3’ end formation are rather virus specific. Of note, the RNA binding protein *TARDBP*/TDP43, whose levels and localization are affected by viral infection, is known to be involved in 3’ end formation, for example by binding close to polyadenylation sites and repressing their usage. It follows that its mis-expression would lead to increased usage of bound polyadenylation sites, while at the same time transcriptional readthrough or other virally induced mechanisms decrease usage of proximal termination sites. On the other hand, *TARDBP*/TDP43 can also favor proximal isoform usage in certain genes (50). In these cases, we would expect a stronger shift towards distal polyadenylation sites upon viral infection, which was indeed reflected in our data. There is now an increasing body of evidence showing that a specific 3’ untranslated region isoform can contribute to pathology related phenotypes. As an example, the extended 3′ UTR of α-synuclein isoform (aSynL isoform) has been previously shown to impact accumulation of α-synuclein protein, and its relocalization from synaptic terminals to mitochondria, characteristic of Parkinson’s disease pathology. Interestingly, we could also observe an expression of aSynL isoform exclusively in HSV-1 infected organoids, however due to low number of reads covering this region, this observation will require further investigation.

We now discuss one of the perhaps most interesting questions, i.e., do we see indications that HSV-1 infection is possibly involved in triggering neurodegeneration?

Progressive loss of neuronal structure and function can be driven by different cellular alterations. In our system we could observe several molecular changes which can be directly linked to neurodegeneration processes. Firstly, dysregulation of the extracellular matrix (ECM) genes and ECM disruption can lead to synaptic and neuronal loss, promoting the pathogenesis of neuropathological states (59) and neurodegeneration (59). Interestingly, we observed upregulation of ECM enzymes involved in 3-*O*-sulfation of heparan sulphate such as *HS3ST2* and *HS3ST6* in infected organoids. 3-*O*-sulfation of heparan sulphate results not only in dramatic increase of susceptibility to viral infection (60), but also causes heparan sulphate internalization, promoting abnormal phosphorylation of tau in Alzheimer’s disease-related tau pathology (61).

Secondly, we observed specific molecular alterations linked to neurodegeneration: **(a)** downregulation of the microtubule regulator STMN2, that has been previously associated with neurodegenerative diseases (46), **(b)** TDP43 protein downregulation and/or relocalization from the nucleus, which is a hallmark of diseases associated with *TARDBP*/TDP43 proteinopathies such as amyotrophic lateral sclerosis (ALS), frontotemporal dementia (FTD) and Alzheimer’s disease (AD) (25, 49), **(c)** upregulation of the small noncoding RNA BC200, which was shown to be associated with AD and variety of tumors (24, 62). The upregulation of BC200 during HSV-1 infection is most likely associated with widespread induction of antisense transcription.

BC200 is selectively expressed in synaptodendritic neuronal compartments, where it is thought to modulates local protein synthesis and in normal aging, it is reduced by > 60% between the ages of 49 and 86. In contrast, in AD brains BC200 RNA was significantly dysregulated in the brain areas that are involved in the disease (24). However due to contradictory data in the context of AD (63), further investigation will be necessary to elucidate the expression and function of this interesting noncoding RNAs in diseased brains.

Finally, our data indicate that HSV-1 infection very rapidly alters neuronal tissue functionality as indicated by decreased firing rates and loss of coordinated neuronal activity. Extensive studies from human AD patients and animal models suggest that destabilization of spontaneous neuronal activity in cortical and hippocampal circuits is a typical feature of AD pathophysiology and can be detected already at very early stages of the disease (64–66). We suggest that it is crucial for future studies to combine highly resolved electrophysiological data with spatial transcriptomics, in order to define the transcriptome changes accompanying functional defects and their association with neurodegenerative processes.

Taken together, our data uncover several molecular and functional defects of neuronal tissue, directly or indirectly caused by HSV-1 infection. We suggest that further studies should focus on elucidating the exact chain of molecular events (including earlier events than covered in our approach) leading to the observed signatures of neurodegeneration, and how these may be tackled with antiviral or more targeted therapies.

## Methods

### iPSC lines

The human iPSC lines iPSC-1 XMO01 (67) and iPSC-2 (A18945, Thermo Fisher Scientific) were cultured in standard conditions (37°C, 4% CO_2_, and 100% humidity), in E8 Flex medium (Thermo Fisher Scientific).

### Generation of cerebral organoids

We generated iPSC-derived cerebral organoids according to a protocol previously described with some modifications (27, 28). Shortly, after dissociation into single-cell suspension with accutase, we seeded 6,000 cells per one well of 96-well plates in 100 µl of embryoid body (containing: DMEM/F12, 20% Knockout replacement serum, 1x Glutamax, 1x MEM-NEAA, 2% ESC FBS, 50µM ROCK Inhibitor, 10 µM bFGF) medium. After 4 days, we replaced the medium with EB medium without bFGF and ROCK inhibitor. On day 6, we replaced the medium with a neural induction medium (NIM: DMEM/F12, 1x N2 supplement, 1x Glutamax, 1x MEM-NEAA, 10µg/ml heparin solution). At day 7-9, we embedded the formed organoids into Matrigel (Corning, 356234) and kept them in NIM for two days, and in organoid differentiation medium containing 1:1 DMEM/F12: Neurobasal, 1xN2 supplement, 1x B27-vitamin A supplement, insulin, 2-ME solution, Glutamax, MEM-NEAA, CHIR99021 for another four days. Next, we transferred the organoids to ultra-low attachment 6-well plates and cultured them on an orbital shaker (80 rpm) in organoid maturation medium containing 1:1 DMEM/F12: Neurobasal, N2 supplement, B27+ vitamin A supplement, insulin, 2-ME solution, Glutamax supplement, MEM-NEAA, Sodium Bicarbonate, Vitamin C solution, chemically defined lipid concentrate, BDNF, GDNF, cAMP, 1% Matrigel.

### Viral infections

The following HSV-1 strains were employed in this study: strain 17 containing GFP under an MCMV promoter (68) KOS (VR-1493; ATCC), and a KOS-based ICP27 knockout virus (52). Around 5-10 organoids cultured in 6-well plates were infected with 75,000 PFU of HSV-1/ per organoid (corresponding to an MOI of 1 as calculated from the estimated number of cells at the surface of the organoids), and 24 h after the medium was replaced with fresh organoid maturation medium and organoids were cultured for an additional 3 or 6 days (3 dpi/6 dpi). The control organoids were culture in parallel in the same conditions, but without viral infection.

### PCR analyses and Nanostring

We performed gene expression analysis by quantitative real-time RT-PCR (qPCR) using SYBR Green PCR Master Mix (Thermo Fisher Scientifics, #4309155) and the ViiA™ 7 Real-Time PCR System (Applied Biosystems). For each target gene, we measured cDNA samples and negative controls in triplicates using ABI PRISM™ 384-well Clear Optical Reaction Plate (Applied Biosystems/ Thermo Fisher Scientifics, #4309849). We calculated the relative transcript levels of each gene based on the 2− ΔΔCT method or as the percentage of housekeeping gene expression. All primer sequences are reported in Table S3. We performed Nanostring-based differential expression analyses of mRNA expression using a custom-designed 72-plex Nanostring nCounterTM probes panel. The mRNA transcript quantification analysis, including sample preparation protocol, hybridization, and detection was performed according to manufacturer’s recommendations.

### Immunostaining of brain organoids

For immunostaining organoids were washed three times with PBS and fixed tissue in 4% paraformaldehyde for 20 to 60 min (depending on the organoid size) at 4 °C, washed with PBS three times 10 min. The tissue was incubated in 40% sucrose (in PBS) till it sunk (overnight) and embedded in 13%/10% gelatin/sucrose. Frozen blocks were stored at −80 °C, prior cryosection. 10-14 μm sections were prepared using cryostat (Leica). Sections were incubated with warm PBS 10-15 min, to remove the embedding medium and next fixed for additional 10 min with 4% PFA, washed three times with PBS and blocked and permeabilized in 0.25% Triton-X, 5% normal goat serum in PBS for 1 h. Sections were incubated with primary antibodies (Table S3) in 0.1% Triton-X, 5% normal goat serum overnight, and the washed three times 10 min with PBST (0.1% Triton X-100) and incubated with secondary antibodies at RT for 2 h, stain with DAPI (final 1 µg/ml) for 10 min and washed three times with PBST. The images were acquired using a Keyence BZ-X710 (Osaka, Japan) microscope.

### Western Blotting

Organoids were lysed in RIPA buffer (150 mM NaCl, 5 mM EDTA, 50 mM Tris, 1% NP-40, 0.5% sodium deoxycholate and 0.1% SDS) in the presence of protease and phosphatase inhibitors. Subsequently, the samples were incubated on ice for 30 minutes and centrifuged at 14,000 x g for 20 minutes at 4 °C. The total protein concentrations were determined using the Pierce™ BCA assay (Thermo Fisher Scientific, #23225). Samples (10-20 µg) were boiled in standard SDS-PAGE sample buffer supplemented with 100 mM DTT, boiled at 95 °C and resolved in a 10–12% SDS–PAGE. Proteins were transferred onto Transblot Turbo Midi PVDF membranes (Bio-Rad, #1704157). The membrane was blocked with Tris-buffered saline containing 0.1% Tween-20 (TBST, Sigma-Aldrich) supplemented with 5% skimmed milk for 1 h at room temperature. Subsequently, the membranes were incubated with the primary antibodies (Table S3) diluted in TBST with skimmed milk at 4 °C overnight. After washing in TBST, the membranes were incubated with anti-rabbit or anti-mouse horseradish peroxidase-conjugated secondary antibody (Cell Signaling, Frankfurt/Main, Germany) for 1 h at room temperature. Blots were then developed with AmershamTM ECLTM select reagent (Cytiva, RPN3243) and visualized by Fusion FX (Vilber, Collégien, France).

### RNA-sequencing

For the total RNA sequencing we used 100 ng of total RNA, where rRNA was depleted using RNase H-based protocol. We mixed total RNA with 1 μg of a DNA oligonucleotide pool comprising 50-nt long oligonucleotide mix covering the reverse complement of the entire length of each human rRNA (28S rRNA, 18S rRNA, 16S rRNA, 5.8S rRNA, 5S rRNA, 12S rRNA), incubated with 1U of RNase H (Hybridase Thermostable RNase H, Epicentre), purified using RNA Cleanup XP beads (Agencourt), DNase treated using TURBO DNase rigorous treatment protocol (Thermo Fisher Scientific), and purified again with RNA Cleanup XP beads. We fragmented the rRNA-depleted RNA samples and processed them into strand-specific cDNA libraries using TruSeq Stranded Total LT Sample Prep Kit (Illumina) and then sequenced them on NextSeq 500, High Output Kit, 2 x 76 cycles. We performed differential gene expression analysis using the DESeq2 (version 1.20.00) R package.

### RNA-seq data analysis

RNA-seq reads were mapped to the reference containing human GRCh38 and HSV-1:GFP genomes using STAR version 2.7.1a (69). Default settings were used, with the exception of ‘--outFilterMismatchNoverLmax 0.05’. Reads were counted using htseq-count (version 0.9.1 (70)) with GENCODE v27 genome annotation reference (71). Principal component analysis was done on raw read counts after performing variance stabilizing transformation using the vst function from the DESeq2 package version 1.26.0 (72).

### Differential gene expression analysis

Differential gene expression analysis was done using DESeq2 (version 1.26.0), using default options, and multi-factor design that measures the effect of infection while accounting for differences between cell lines (design = ∼ cell_line + day_post_infection). Significance threshold was set to adjusted P-value of 0.05 and log2-fold-change of 1. Gene ontology enrichment analysis was done using the gprofiler2 R package (73). All genes expressed in our system were used as background. GO terms with adjusted p-value lower than 0.05 were considered significantly enriched.

### Antisense transcription analysis

Antisense transcribed regions were identified and defined from STAR-aligned reads using the running sum algorithm described (31). Overlapping antisense transcripts from different RNA-seq libraries were collapsed together. All putative antisense transcripts with an annotated upstream sense gene within 1 kb were discarded as potential readthroughs, and those with a downstream sense gene within 10 kb were discarded as potential transcripts from an upstream TSS. We extended the GENCODE v27 genome annotation reference with detected antisense transcripts and counted the mapping reads using htseq-count 0.9.1, as described earlier.

### FLAM-seq library preparation and data analysis

FLAM-seq libraries were prepared as described (23), starting with 2 micrograms of total RNA for three replicates of non-infected and three replicates of infected (3 dpi) 60 days-old organoids. A detailed version of the protocol can be found on *protocolexchange* (10.21203/rs.2.10045/v1). The only modifications in the protocol were the incubation of the RT reaction for 90 minutes at 42 °C, and the PCR amplification of cDNA performed for 23 cycles with an extension time of 5 minutes per cycle. Libraries were then submitted to the BIMSB Genomics core facility for PacBio amplicon template preparation (SMRTbell Express Template Prep Kit 2.0, 100-938-900) and sequencing on the PacBio Sequel (Sequel Binding and Internal Ctrl Kit 3.0, 101-626-600). Sequencing was performed on diffusion mode on two SMRTcells for each sample and then raw data were preprocessed with SMRTlink v8.0.0.80.529 for CCS generation with a minimum number of 3 passes and a minimum predicted accuracy of 0.99. The CCS reads were then used as input to the FLAM-seq analysis pipeline (23) that outputs gene assignment and poly(A) tail length and sequence for each read. These data, deposited along with the CCS reads on GEO (74) were used for the plots showing read counts and poly(A) tail lengths in Figures 3, **Figure S3** and **Table S1**. As a reference genome and genome annotation, we used GRCh38 with the addition of the HSV-1 genome (75) and Gencode v27 respectively.

Statistical testing of differences in poly(A) tail length between non-infected and infected samples was performed on the merged 3 replicates per condition with the same method and code as described in detail (23). Briefly, the error of tail length measurement for each transcript was interpolated by fitting a linear model to the standard deviations observed using synthetic spike-ins of known lengths. For any given sequenced tail, a new length was sampled from a normal distribution given the aforementioned expected standard deviation, and a p-value was then computed by comparing the measured median tail length with the median tail length of the distribution sampled 1000 times. An additional cutoff at 25 nt was set to increase specificity.

Gene ontology terms enrichment analysis was performed with Gorilla (76), by using genes with significant poly(A) tail length difference between non-infected and infected samples as input (separately for genes with longer or shorter tails in one condition versus the other), and all detected genes in the FLAM-seq data as background. The GO terms “Positive Regulation of Viral Transcription” and “Regulation of Viral Transcritption”, driven by the genes GTF21, DHX9, CDK9, POLR2K, NELFCD, NUCKS1, NELFE and CCL5, were the only ones found enriched with an adjusted p-value lower than 0.05.

Three prime end annotation was performed as in (23) on the merged replicates for infected and uninfected organoids by peak detection on aligned reads 3’ end positions with the Python peakutils module (https://bitbucket.org/lucashnegri/peakutils), by setting a minimum distance of 30 nt between the peaks, keeping only genes with at least 5 read counts, and keeping only peaks supported by at least 2 reads. Reads from the two conditions (infected and uninfected) were then assigned to the detected peaks by simply ascribing the 3’ end position of the read alignment to the closest peak within a 15 nt distance. An isoform index was computed by sorting isoforms for each gene by coordinate: the most 5’ isoform on the gene strand was annotated with an index of 1, which increases distally until the last isoform for that given gene is detected. For the two conditions, isoform usage was computed as the fraction of read counts for a given isoform on the read counts for a given gene. To increase accuracy in isoform usage estimates, only genes with at least 10 read counts were retained when comparing usage between conditions (**Table S2**).

High readthrough genes shown in Figure 3D were annotated as those genes having more than 25% readthrough at 8 hours post infection (WT virus) in (38).

*TARDBP*/TDP43 binding for each isoform was obtained by downloading iCLIP data from *imaps.genialis.com*. Bed files of CLIP sites from the entries 72094, 72102, 69197, 65156, 69164, 69172, 69148, 69124, 69132, 69098, 69106, 69114, 69090, 69074, 69082, obtained from a variety of human cell lines and human samples, were merged and intersected with bedtools to get CLIP sites overlapping a sequence of 40 nt around FLAM-seq-annotated 3’ ends. Functional annotation of *TARDBP*/TDP43-regulated isoforms was retrieved from Rot et al. (2017) where we collected the list of genes in which *TARDBP*/TDP43 either significantly promoted proximal isoform usage or inhibited distal isoform usage.

Statistical testing for the difference in distributions of tail lengths and isoform usage was performed with the R implementation of the Wilcoxon-Mann-Whitney test. This was applied to compare proximal (isoform index = 1) and distal (isoform index > 1) isoform usage in non-infected and infected organoids, as well as to compare the difference in proximal isoform usage between uninfected and infected samples in genes with high readthrough versus the rest of the genes, and in isoforms stratified by *TARDBP*/TDP43 binding or regulation.

The FLAM-seq reads and poly(A) tail visualization were produced by generating a track hub on the BIMSB local copy of the UCSC genome browser. Reads alignments (bam files) resulting from the FLAManalysis pipeline were appended with the tail sequence trimmed by the FLAManalysis pipeline and are therefore visible on the genome browser as a stretch of mismatches in red.

### Single cell RNA-seq

Methanol-fixed cells were centrifuged at 3000–5000 × g for 5 min, rehydrated in 1 ml PBS + 0.01% BSA supplemented with RNAse inhibitors (1 unit/μl RiboLock, ThermoFisher), pelleted and resuspended again in 0.5 ml PBS + 0.01% BSA in the presence of RNAse inhibitors. Cells were manually counted by means of a hemocytometer and diluted to a suspension of typically ∼300 cells/μl in PBS + 0.01% BSA. Cells were encapsulated together with barcoded microparticles (Macosko-2011-10 (V+), ChemGenes Corp. Wilmington, MA, USA) using the Dolomite Bio Nadia instrument, using the standard manufacturer’s dropseq-based (Macosko et al. 2015) scRNA-seq protocol.

Droplets were broken immediately after collection and cDNA libraries generated as previously described (https://doi.org/10.1016/ j.isci.2021.102151). First strand cDNA was amplified by equally distributing beads from one run to 24 Smart PCR reactions (50 μl volume; 4 + 9 to 11 cycles). 20 μl fractions of each PCR reaction were pooled (total = 480 μl), then double-purified with 0.6 x volumes of AMPure XP beads (Beckman Coulter). Amplified cDNA libraries were assessed and quantified on a BioAnalyzer High Sensitivity Chip (Agilent) and the Qubit dsDNA HS Assay system (ThermoFisher). 600 pg of each cDNA library was fragmented, amplified (13 cycles) and indexed for sequencing with the Nextera XT v2 DNA sample preparation kit (Illumina) using custom primers enabling 3’-targeted amplification. The libraries were purified with AMPure XP Beads, quantified and sequenced on Illumina NextSeq500 sequencers (library concentration 1.8 pM; NextSeq 500/550 High Output v2 kit (75 cycles) in paired-end mode; read 1 = 20 bp using the custom primer Read1CustSeqB (39) read 2 = 64 bp).

## Single-cell RNA-seq data analysis

### Quality control and data pre-processing

Single-cell gene read count tables were generated using the PiGx-scRNA-seq pipeline (https://doi.org/10.1093/gigascience/giy123) version 1.1.4. Data was analyzed in R (v3.6) using Seurat (v3). Cells with less than 500 detected genes were filtered out, gene expression was normalized using the SCTransform workflow and the different samples (control, 3d pi, 6 dpi) were then integrated using a mutual nearest neighbor approach as illustrated on the Seurat website (77). Dimensionality reduction was performed using PCA and we selected 30 PCs based on Elbow plot. The FindClusters function which implements shared nearest neighbor (SNN) modularity optimization-based clustering algorithm was applied with a resolution of 1.5 and identified in 45 initial clusters for the entire dataset, and 21 for only ctrl samples. The UMAP algorithm was used to create two-dimensional projections to visualize the data.

For identifying the markers for each cluster, the “FindAllMarkers” function with default parameters was used, identifying negative and positive markers for that cluster. To label clusters, we performed a comprehensive manual annotation of the clusters.

### Local density plot

The relative density between uninfected and HSV infected organoids on UMAP was measured using 2D kernel density estimation (MASS v.7.3-53) by each experiment (78). log2 ratio between estimated density was projected on UMAP.

### Cell proportion

Differences in cell type proportions were analyzed using mixed-effects linear models (lme4 v.1.1-23) using a binomial model with cell type as fixed effect and replicates as random effect (73) p-value less than 0.001 was considered statistical significant.

### Differential gene expression and gene set enrichment analysis

In order to identify differentially regulated genes among experiments, using Wilcoxon rank sum test by scTransform normalized counts was utilized. The adjusted p-value was calculated using Bonferroni Correction for multiple testing correction.

To identify enriched pathways associated with HSV infection, GSEA using the fgsea (v1.10.1) was performed with 10,000 permutation. Ranking metric was done by logFC from Differential expression analysis. Two databases (REACTOME, HALLMARK pathway from the MsigDB) were used as reference pathways. Only pathways with with more than 3 statistical significant reference genes, FDR < 0.05 were considered as significantly enriched.

### Tissue processing and Visium data generation

Brain organoid samples, frozen in isopentane and embedded in OCT (TissueTek Sakura) and cryosectioned at −12 °C (Thermo Cryostar). Sections were placed on chilled Visium Spatial Gene Expression Slides (2000233, 10X Genomics). Tissue sections were then fixed in chilled methanol and stained according to the Visium Spatial Gene Expression User Guide (CG000239 Rev A, 10X Genomics).

Libraries were prepared according to the Visium Spatial Gene Expression User Guide (CG000239, https://assets.ctfassets.net/an68im79xiti/3pyXucRaiKWcscXy3cmRHL/a1ba41c 77cbf60366202805ead8f64d7/CG000239_VisiumSpatialGeneExpression_UserGuide_Rev_A.pdf). Libraries were sequenced on a NovaSeq 6000 System (Illumina) using a NovaSeq S4 Reagent Kit (200 cycles, 20027466, Illumina), at a sequencing depth of approximately 250-400M read-pairs per sample. Sequencing of 10 µm organoid slices (four organoids/condition) yielded, for uninfected organoids, on average 8,000 unique molecular identifiers (UMIs), which quantified ∼3,700 genes per spot.

### Label transfer from scRNA to spatial transcriptomic datasets

We transferred labels from each of the clusters identified in the scRNAseq dataset to each spot in the spatial transcriptomic dataset using Seurat V3 (ref: Comprehensive Integration of Single-Cell Data (77). Briefly, we used the SelectIntegrationFeatures with default parameters to identify the gene to use during the label transfer process and the FindTransferAnchors function (parameters: normalization.method = “SCT”, reference.assay = ‘SCT’, query.assay = ‘SCT’, reduction = ‘cca’, dims = 1:30) to find the set of anchors between the single cell and the query spatial transcriptomic datasets. We then used the TransferData function (parameters: prediction.assay = TRUE, weight.reduction = ‘cca’, dims = 1:30) to obtain the prediction scores of each spatial transcriptomic spot for each of the single cell cluster.

### Multi-electrode array (MEA) recording

Spontaneous extracellular field potential in uninfected and infected 60 days old organoids generated from iPSC line-2 were acquired in 6-well MEA plates with 64 poly-3,4-ethylendioxythiophen (PEDOT) electrodes (Axion Biosystems, Atlanta, GA, USA). Each well was previously coated with 100 μg/mL poly-D-lysine. In the following, 60 days old organoids were placed in the middle of each well. Subsequently, Geltrex^TM^ was added on top of the organoid and placed for 30 min in the incubator to spatially fix the organoid. Afterwards, BrainPhys^TM^ media supplemented with N2 supplement, B27+ vitamin A supplement, 20 ng/ml BDNF, 20 ng/ml GDNF, 0.5 mM dibutyryl cyclic-AMP and 200 nM ascorbic acid was added to the wells. Two days after organoid placement, recording was performed using a Maestro MEA system and AxIS Software Spontaneous Neural Configuration (Axion Biosystems). Spikes were detected with AxIS software using an adaptive threshold crossing set to 6 times the standard deviation of the estimated noise for each electrode (channel). The plate was first allowed to rest for three minutes in the Maestro device, and then 5 minutes of data were recorded. For the MEA analysis, the electrodes that detected at least 5 spikes/min were classified as active electrodes using Axion Biosystems’ Neural Metrics Tool. One hour after recording, three of six organoids were infected with 75,000 PFU of HSV-1 GFP/ per organoid and recordings were performed every hour for 48 h after 1 h of infection. We included in the analysis only the electrodes that were classified as active electrodes at the time 1 h before infection. In every well, the ‘timepoint median’ was obtained by computing the median value of the number of spikes across active electrodes. For the heatmap in Figure 2D, we ranked each electrode in a single well by the total number of spikes measured and selected the four electrodes with the highest total number of spikes. In order to have a dynamic range between 0 and 1, the number of spikes was max-normalized. Briefly, the number of spikes measured across all electrodes and timepoints in a single sample was divided by the maximum number of spikes measured in that sample.

### Calcium Imaging

For calcium imaging experiments, organoids were plated in 35 mm glass bottom imaging dish coated with poly-D-lysine and Geltrex^TM^. Two days after plating, organoids were loaded with 5μM CalBryte590 AM (AAT Bioquest) and 0.02% Pluronic acid in organoid maturation media for 30 min at 37 °C. Imaging was performed in a Dragonfly spinning disk confocal microscope (Andor, Oxford Instruments) at a frequency of 5 Hz for 5 min. The resulting images were then background subtracted and the relative changes in intensity over time were computed. Activity-based segmentation was performed as in Junek at al. 2009 (79). Spike detection was performed on the intensity traces from each ROI to calculate AP firing frequency.

## Acknowledgments

The authors declare no competing financial or commercial interests. We thank Maryna Korshevniuk for the initial data analysis, Ruth Pareja, for the technical support with iPSC culture and western blotting, Anastasiya Boltengagen and Salah Ayoub for the technical support with single-cell RNA-seq. Margareta Herzog for the lab organizational support. We thank Claudia Quedenau, Ali Kerim Secener, Tatiana Borodina (NGS Unit) and Thomas Conrad (Single Cell Technologies) from the BIMSB Genomics platform for performing sequencing runs and preprocessing of sequencing data of VISIUM and PacBio. We thank Jonathan Alles for his fundamental help with 3’ end annotation analysis. We thank Sebastian Diecke (BIH Stem Cell facility) for providing and biobanking iPSC lines, Heiko Lickert for providing the XMO01 iPS cell line. We thank Anna Cliffe, Lars Dölken, Adam Whisnant and Thomas Hennig for fruitful discussions and support. We also thank all the members of the N. Rajewsky laboratory for critical and useful discussions. This work was supported by BIH Grant: Multiscale Omics Projects. TMP was a member of the graduate school Berlin School of Integrative Oncology (BSIO) and partially funded by Deutsche Akademische Austauschdienst (DAAD).

## Data deposition

The GEO identifier: GSE163952

## Author Contributions

Conceptualization: A.R.-W., E.W., N.R., Methodology: A.R.-W., E.W., A.L., Calcium based Imaging: A.O.-M., A.W. Computational analysis: E.W., S.J.K., T.M.P., B.B, - single-cell RNA-seq, I.L. - FLAM-seq, P.G. – bulk RNAseq, E.W., S.J.K. and T.M- spatial VISIUM. Writing – Original Draft: A.R.-W. Writing – Review & Editing: I.L., E.W., A.L., P.G., M.L, and NR. All Authors discussed and analyzed the data. Supervision: N.R., M.L.

## Conflict of interest

The authors declare that they have no conflict of interest.

## Notes

### Competing Interest Statement

The authors have declared no competing interest.

