## Supplementary_Figures for "Neurodegeneration in human brain organoids infected with herpes simplex virus type 1"

#### Figure S1.

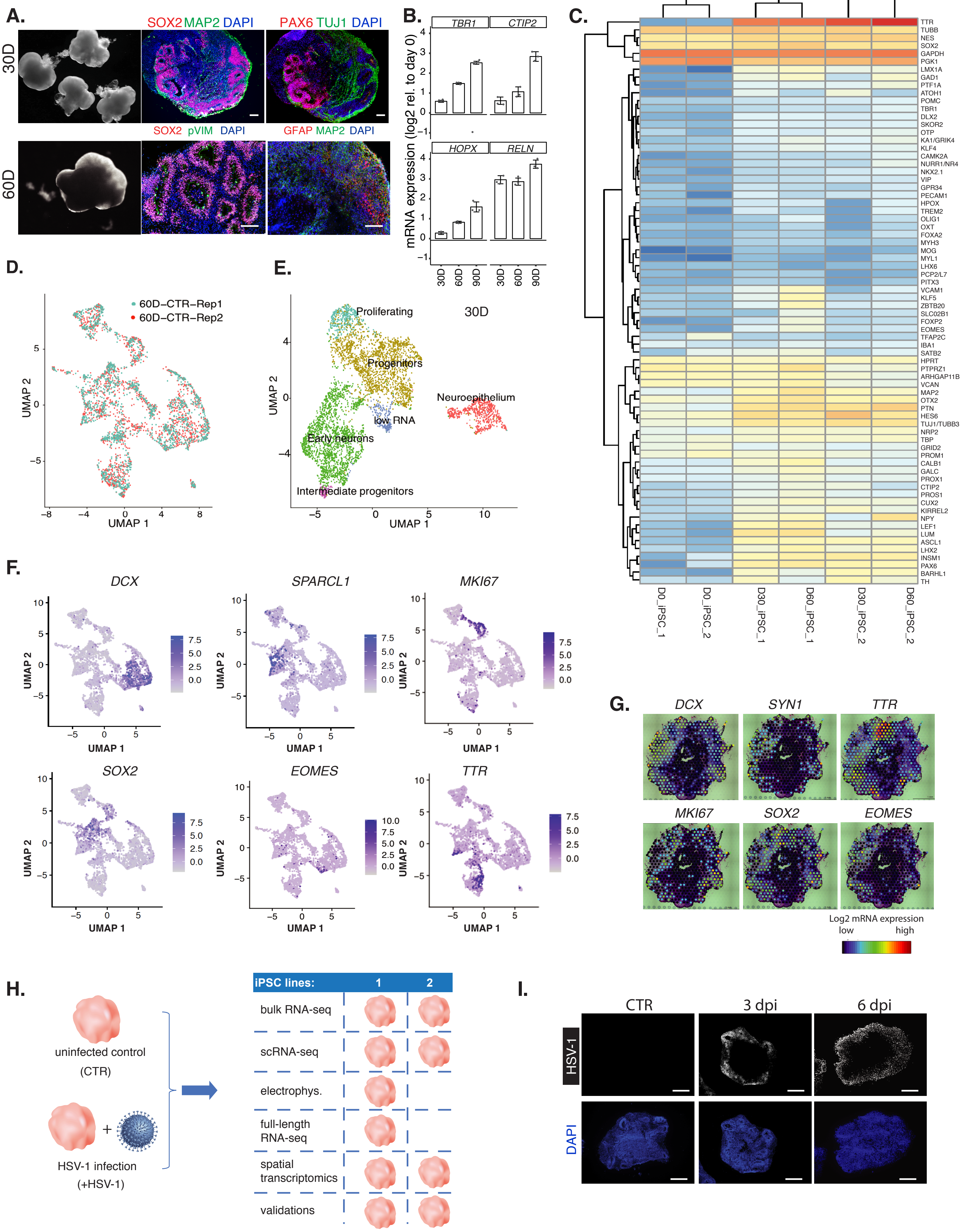
A.
B.
C.
D.
E.
F.
G.
H.
I.
30D
60D
SOX2 MAP2 DAPI
PAX6 TUJ1 DAPI
SOX2 pVIM DAPI
GFAP MAP2 DAPI
TBR1
CTIP2
HOPX
RELN
mRNA expression (log2 rel. to day 0)
60D–CTR–Rep1
60D–CTR–Rep2
UMAP 1
UMAP 2
Proliferating
Progenitors
Neuroepithelium
low RNA
Early neurons
Intermediate progenitors
DCX
SPARCL1
MKI67
SOX2
EOMES
TTR
SYN1
Log2 mRNA expression
low
high
D0_iPSC_1
D0_iPSC_2
D30_iPSC_1
D60_iPSC_1
D30_iPSC_2
D60_iPSC_2
uninfected control
(CTR)
HSV-1 infection
(+HSV-1)
iPSC lines:
1
2
bulk RNA-seq
scRNA-seq
electrophys.
full-length RNA-seq
spatial transcriptomics
validations
CTR
3 dpi
6 dpi
HSV-1
DAPI

#### Figure S2

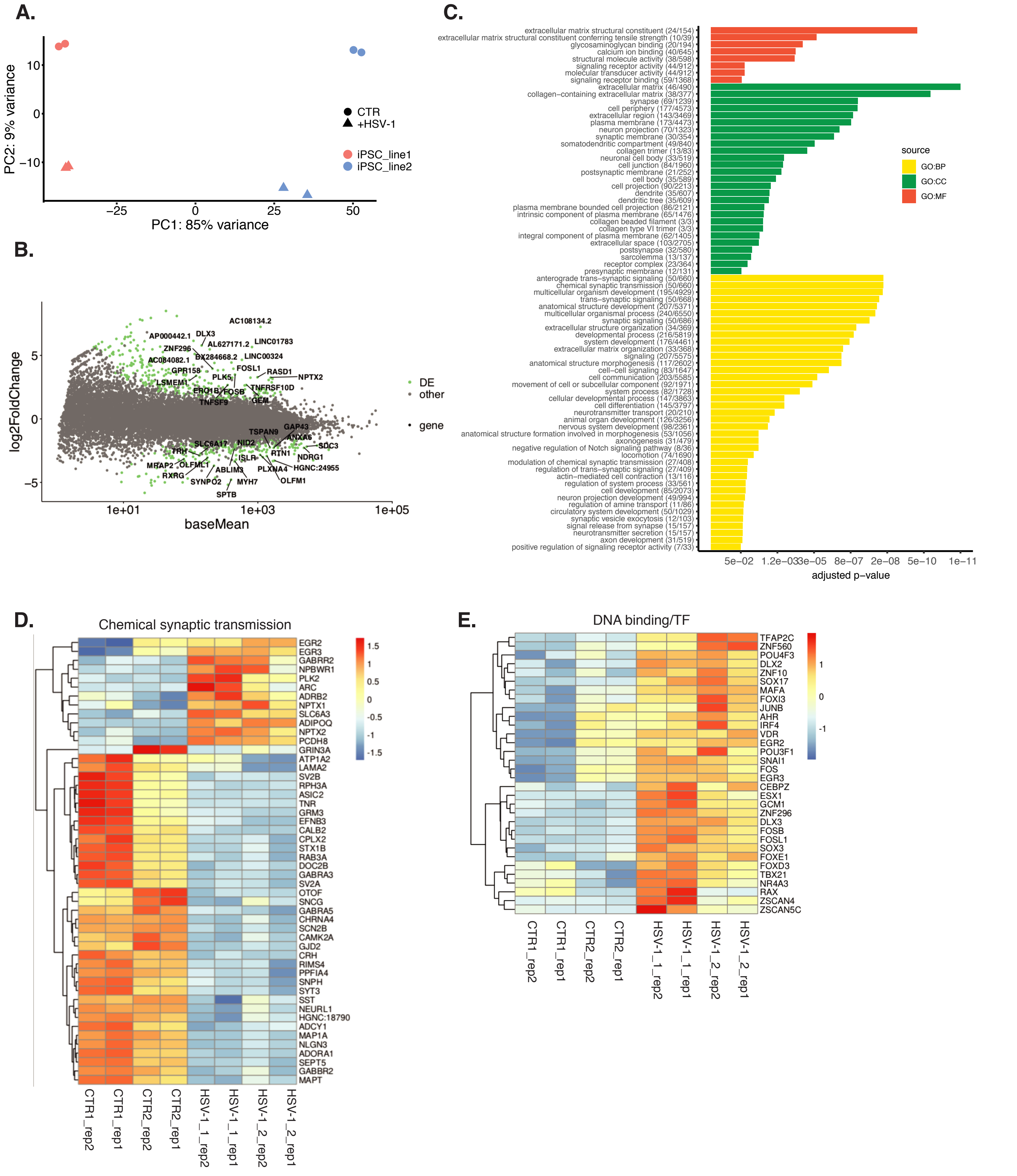
A.
PC2: 9% variance
PC1: 85% variance
CTR
+HSV-1
iPSC_line1
iPSC_line2
B.
log2FoldChange
baseMean
DE
other
gene
C.
adjusted p-value
source
GO:BP
GO:CC
GO:MF
extracellular matrix structural constituent (24/154)
extracellular matrix structural constituent conferring tensile strength (10/39)
glycosaminoglycan binding (20/194)
calcium ion binding (40/645)
structural molecule activity (38/598)
signaling receptor activity (44/912)
molecular transducer activity (44/912)
signaling receptor binding (59/1368)
extracellular matrix (46/490)
collagen-containing extracellular matrix (38/377)
synapse (69/1239)
cell periphery (177/4573)
extracellular region (143/3469)
plasma membrane (173/4473)
neuron projection (70/1323)
synaptic membrane (30/354)
somatodendritic compartment (49/840)
collagen trimer (13/83)
neuronal cell body (33/519)
cell junction (84/1960)
postsynaptic membrane (21/252)
cell body (35/589)
cell projection (90/2213)
dendrite (35/607)
dendritic tree (35/609)
plasma membrane bounded cell projection (86/2121)
intrinsic component of plasma membrane (65/1476)
collagen beaded filament (3/3)
collagen type VI trimer (3/3)
integral component of plasma membrane (62/1405)
extracellular space (103/2705)
postsynapse (32/580)
sarcolemma (13/137)
receptor complex (23/364)
presynaptic membrane (12/131)
anterograde trans-synaptic signaling (50/660)
chemical synaptic transmission (50/660)
multicellular organism development (195/4929)
trans-synaptic signaling (50/668)
anatomical structure development (207/5371)
multicellular organismal process (240/6550)
synaptic signaling (50/686)
extracellular structure organization (34/369)
developmental process (216/5819)
system development (176/4461)
extracellular matrix organization (33/368)
signaling (207/5575)
anatomical structure morphogenesis (117/2602)
cell-cell signaling (83/1647)
cell communication (203/5585)
movement of cell or subcellular component (92/1971)
system process (82/1728)
cellular developmental process (147/3863)
cell differentiation (145/3797)
neurotransmitter transport (20/210)
animal organ development (126/3256)
nervous system development (98/2361)
anatomical structure formation involved in morphogenesis (53/1056)
axonogenesis (31/479)
negative regulation of Notch signaling pathway (8/36)
locomotion (74/1690)
modulation of chemical synaptic transmission (27/408)
regulation of trans-synaptic signaling (27/409)
actin-mediated cell contraction (13/116)
regulation of system process (33/561)
cell development (85/2073)
neuron projection development (49/994)
regulation of amine transport (11/86)
circulatory system development (50/1029)
synaptic vesicle exocytosis (12/103)
signal release from synapse (15/157)
neurotransmitter secretion (15/157)
axon development (31/519)
positive regulation of signaling receptor activity (7/33)
D.
Chemical synaptic transmission
CTR1_rep2
CTR1_rep1
CTR2_rep2
CTR2_rep1
HSV-1_1_rep2
HSV-1_1_rep1
HSV-1_2_rep2
HSV-1_2_rep1
E.
DNA binding/TF

### Figure S2

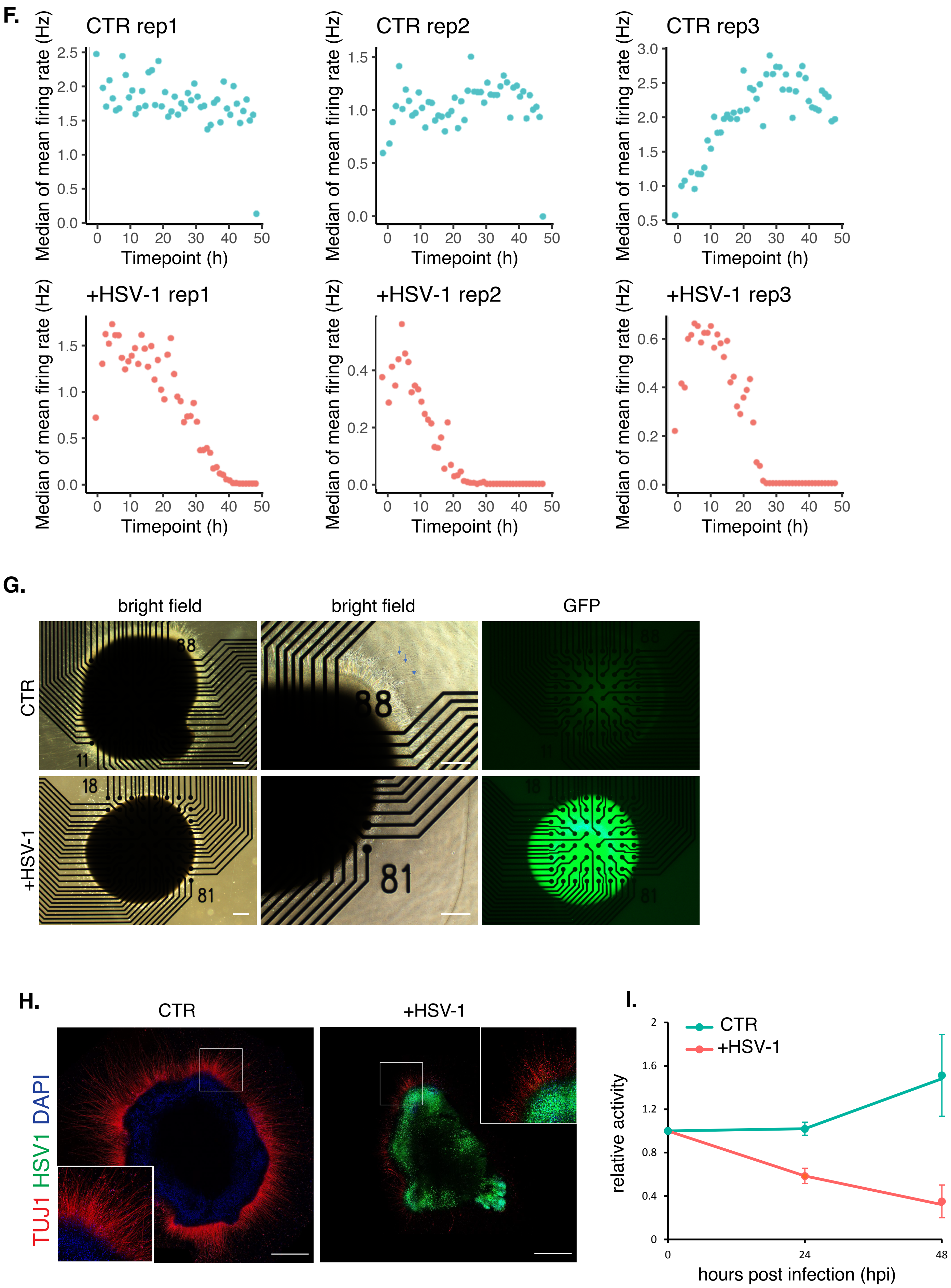
F.
CTR rep1
CTR rep2
CTR rep3
+HSV-1 rep1
+HSV-1 rep2
+HSV-1 rep3
Median of mean firing rate (Hz)
Timepoint (h)
G.
bright field
bright field
GFP
CTR
+HSV-1
88
11
18
81
H.
CTR
+HSV-1
TUJ1 HSV1 DAPI
I.
CTR
+HSV-1
relative activity
hours post infection (hpi)
0
24
48

#### Figure S3

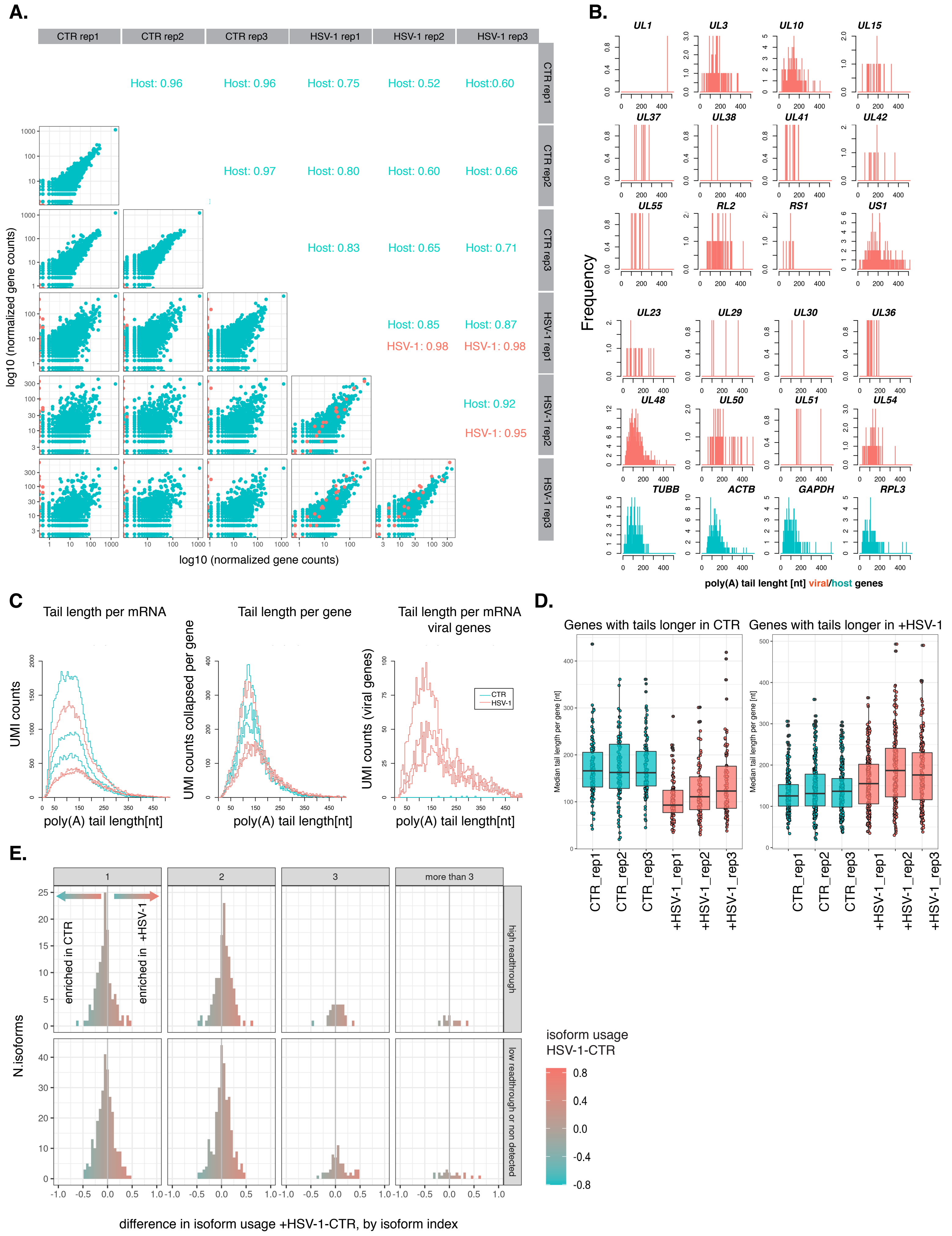

A.
CTR rep1
CTR rep2
CTR rep3
HSV-1 rep1
HSV-1 rep2
HSV-1 rep3
Host: 0.96
Host: 0.96
Host: 0.75
Host: 0.52
Host:0.60
Host: 0.97
Host: 0.80
Host: 0.60
Host: 0.66
Host: 0.83
Host: 0.65
Host: 0.71
Host: 0.85
Host: 0.87
HSV-1: 0.98
HSV-1: 0.98
Host: 0.92
HSV-1: 0.95
log10 (normalized gene counts)
log10 (normalized gene counts)
B.
UL1
UL3
UL10
UL15
UL37
UL38
UL41
UL42
UL55
RL2
RS1
US1
UL23
UL29
UL30
UL36
UL48
UL50
UL51
UL54
TUBB
ACTB
GAPDH
RPL3
Frequency
poly(A) tail lenght [nt] viral/host genes
C.
Tail length per mRNA
Tail length per gene
Tail length per mRNA viral genes
UMI counts
UMI counts collapsed per gene
UMI counts (viral genes)
CTR
HSV-1
poly(A) tail length[nt]
poly(A) tail length[nt]
poly(A) tail length[nt]
D.
Genes with tails longer in CTR
Genes with tails longer in +HSV-1
Median tail length per gene [nt]
CTR_rep1
CTR_rep2
CTR_rep3
+HSV-1_rep1
+HSV-1_rep2
+HSV-1_rep3
E.
1
2
3
more than 3
enriched in CTR
enriched in +HSV-1
high readthrough
low readthrough or non detected
N.isoforms
isoform usage HSV-1-CTR
0.8
0.4
0.0
-0.4
-0.8
difference in isoform usage +HSV-1-CTR, by isoform index

### Figure S4

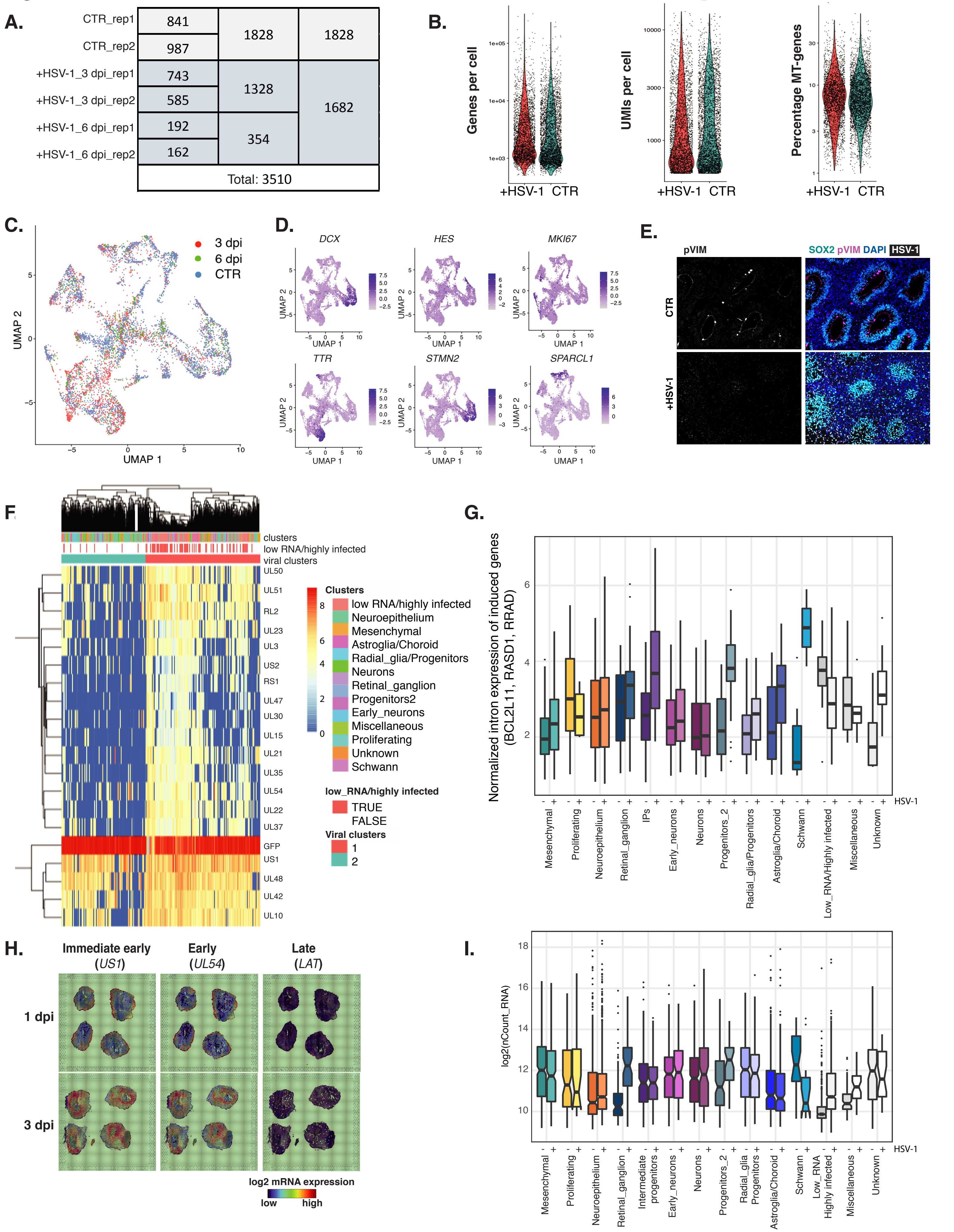
A.
CTR_rep1 841
CTR_rep2 987
1828
1828
+HSV-1_3 dpi_rep1 743
+HSV-1_3 dpi_rep2 585
1328
+HSV-1_6 dpi_rep1 192
+HSV-1_6 dpi_rep2 162
354
1682
Total: 3510
B.
Genes per cell
UMIs per cell
Percentage MT-genes
+HSV-1 CTR
C.
3 dpi
6 dpi
CTR
UMAP 1
UMAP 2
D.
DCX
HES
MKI67
TTR
STMN2
SPARCL1
E.
pVIM
SOX2 pVIM DAPI HSV-1
CTR
+HSV-1
F.
clusters
low RNA/highly infected
viral clusters
UL50
UL51
RL2
UL23
UL3
US2
RS1
UL47
UL30
UL15
UL21
UL35
UL54
UL22
UL37
GFP
US1
UL48
UL42
UL10
Clusters
low RNA/highly infected
Neuroepithelium
Mesenchymal
Astroglia/Choroid
Radial_glia/Progenitors
Neurons
Retinal_ganglion
Progenitors2
Early_neurons
Miscellaneous
Proliferating
Unknown
Schwann
low_RNA/highly infected
TRUE
FALSE
Viral clusters
1
2
G.
Normalized intron expression of induced genes (BCL2L11, RASD1, RRAD)
HSV-1
Mesenchymal
Proliferating
Neuroepithelium
Retinal_ganglion
IPs
Early_neurons
Neurons
Progenitors_2
Radial_glia/Progenitors
Astroglia/Choroid
Schwann
Low_RNA/Highly infected
Miscellaneous
Unknown
H.
Immediate early (US1)
Early (UL54)
Late (LAT)
1 dpi
3 dpi
log2 mRNA expression
low
high
I.
log2(nCount_RNA)
HSV-1
Mesenchymal
Proliferating
Neuroepithelium
Retinal_ganglion
Intermediate progenitors
Early_neurons
Neurons
Progenitors_2
Radial_glia Progenitors
Astroglia/Choroid
Schwann
Low_RNA Highly infected
Miscellaneous
Unknown

#### Figure S4

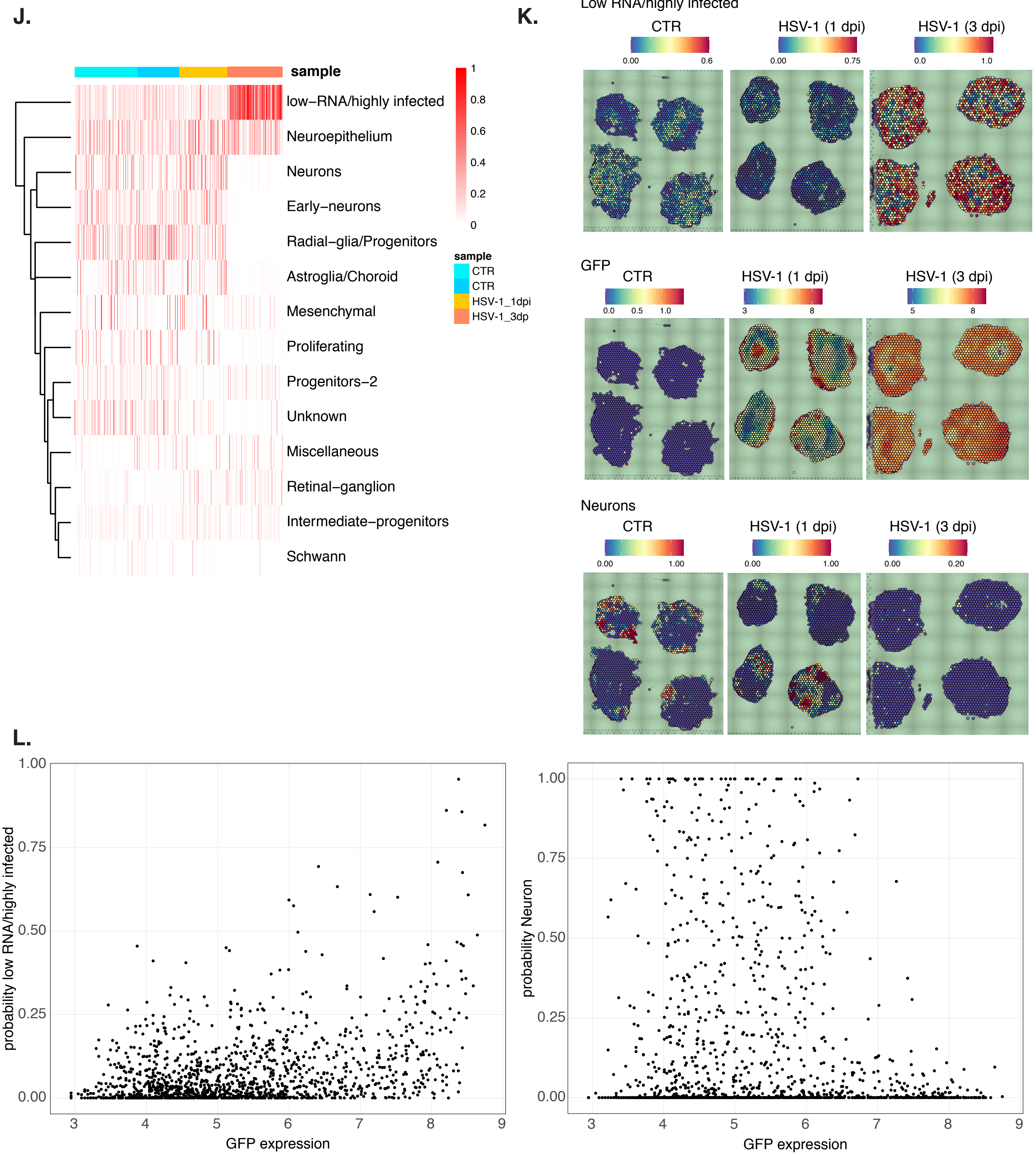

J.
sample
low–RNA/highly infected
Neuroepithelium
Neurons
Early–neurons
Radial–glia/Progenitors
Astroglia/Choroid
Mesenchymal
Proliferating
Progenitors–2
Unknown
Miscellaneous
Retinal–ganglion
Intermediate–progenitors
Schwann
1
0.8
0.6
0.4
0.2
0
sample
CTR
CTR
HSV-1_1dpi
HSV-1_3dp
K.
Low RNA/highly infected
CTR
HSV-1 (1 dpi)
HSV-1 (3 dpi)
0.00
0.6
0.00
0.75
0.0
1.0
GFP
CTR
HSV-1 (1 dpi)
HSV-1 (3 dpi)
0.0
0.5
1.0
3
8
5
8
Neurons
CTR
HSV-1 (1 dpi)
HSV-1 (3 dpi)
0.00
1.00
0.00
1.00
0.00
0.20
L.
probability low RNA/highly infected
1.00
0.75
0.50
0.25
0.00
3
4
5
6
7
8
9
GFP expression
probability Neuron
1.00
0.75
0.50
0.25
0.00
3
4
5
6
7
8
9
GFP expression

### Figure S5

A.

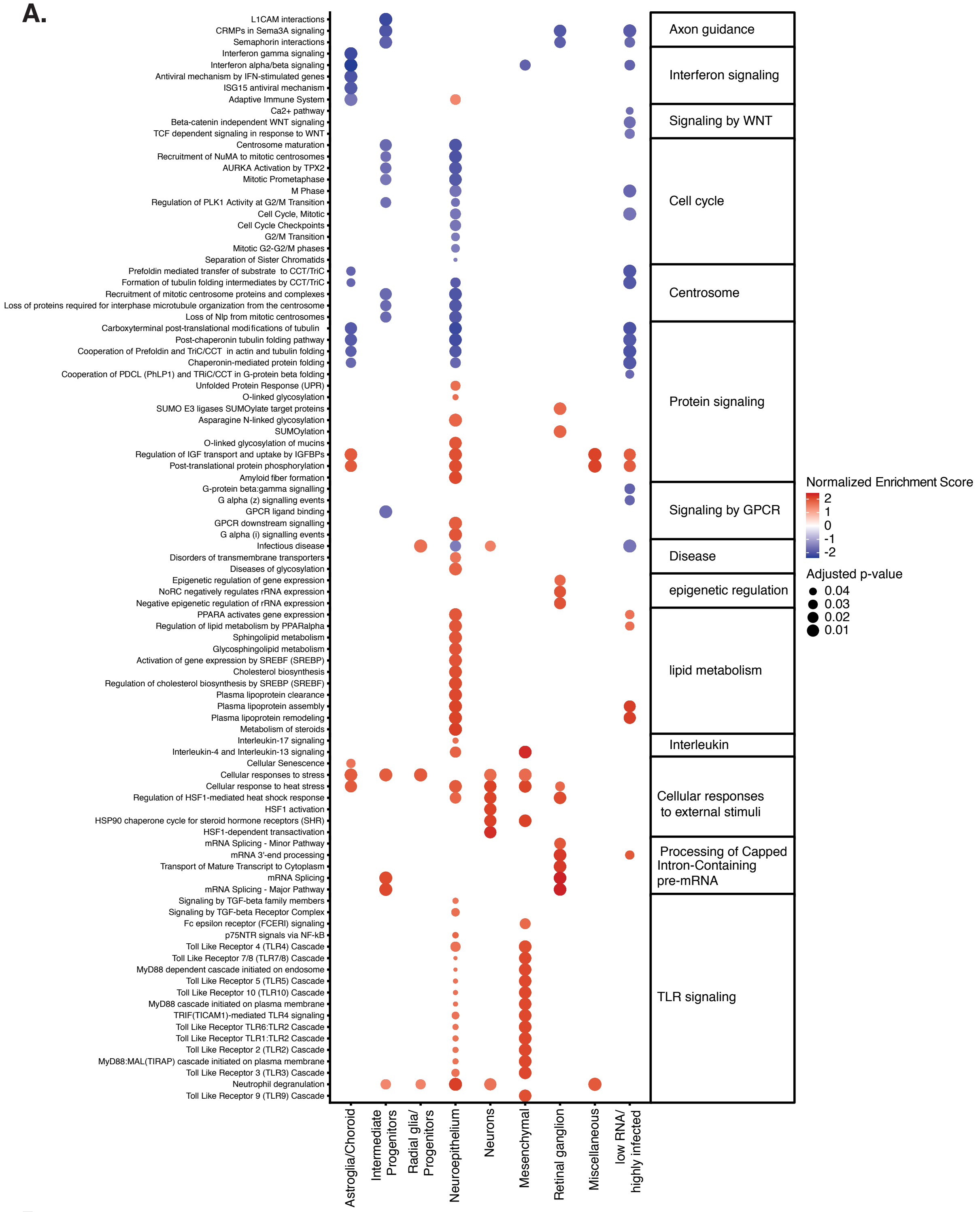

L1CAM interactions
CRMPs in Sema3A signaling
Semaphorin interactions
Interferon gamma signaling
Interferon alpha/beta signaling
Antiviral mechanism by IFN-stimulated genes
ISG15 antiviral mechanism
Adaptive Immune System
Ca2+ pathway
Beta-catenin independent WNT signaling
TCF dependent signaling in response to WNT
Centrosome maturation
Recruitment of NuMA to mitotic centrosomes
AURKA Activation by TPX2
Mitotic Prometaphase
M Phase
Regulation of PLK1 Activity at G2/M Transition
Cell Cycle, Mitotic
Cell Cycle Checkpoints
G2/M Transition
Mitotic G2-G2/M phases
Separation of Sister Chromatids
Prefoldin mediated transfer of substrate to CCT/TriC
Formation of tubulin folding intermediates by CCT/TriC
Recruitment of mitotic centrosome proteins and complexes
Loss of proteins required for interphase microtubule organization from the centrosome
Loss of Nlp from mitotic centrosomes
Carboxyterminal post-translational modifications of tubulin
Post-chaperonin tubulin folding pathway
Cooperation of Prefoldin and TriC/CCT in actin and tubulin folding
Chaperonin-mediated protein folding
Cooperation of PDCL (PhLP1) and TRiC/CCT in G-protein beta folding
Unfolded Protein Response (UPR)
O-linked glycosylation
SUMO E3 ligases SUMOylate target proteins
Asparagine N-linked glycosylation
SUMOylation
O-linked glycosylation of mucins
Regulation of IGF transport and uptake by IGFBPs
Post-translational protein phosphorylation
Amyloid fiber formation
G-protein beta:gamma signalling
G alpha (z) signalling events
GPCR ligand binding
GPCR downstream signalling
G alpha (i) signalling events
Infectious disease
Disorders of transmembrane transporters
Diseases of glycosylation
Epigenetic regulation of gene expression
NoRC negatively regulates rRNA expression
Negative epigenetic regulation of rRNA expression
PPARA activates gene expression
Regulation of lipid metabolism by PPARalpha
Sphingolipid metabolism
Glycosphingolipid metabolism
Activation of gene expression by SREBF (SREBP)
Cholesterol biosynthesis
Regulation of cholesterol biosynthesis by SREBP (SREBF)
Plasma lipoprotein clearance
Plasma lipoprotein assembly
Plasma lipoprotein remodeling
Metabolism of steroids
Interleukin-17 signaling
Interleukin-4 and Interleukin-13 signaling
Cellular Senescence
Cellular responses to stress
Cellular response to heat stress
Regulation of HSF1-mediated heat shock response
HSF1 activation
HSP90 chaperone cycle for steroid hormone receptors (SHR)
HSF1-dependent transactivation
mRNA Splicing - Minor Pathway
mRNA 3'-end processing
Transport of Mature Transcript to Cytoplasm
mRNA Splicing
mRNA Splicing - Major Pathway
Signaling by TGF-beta family members
Signaling by TGF-beta Receptor Complex
Fc epsilon receptor (FCERI) signaling
p75NTR signals via NF-kB
Toll Like Receptor 4 (TLR4) Cascade
Toll Like Receptor 7/8 (TLR7/8) Cascade
MyD88 dependent cascade initiated on endosome
Toll Like Receptor 5 (TLR5) Cascade
Toll Like Receptor 10 (TLR10) Cascade
MyD88 cascade initiated on plasma membrane
TRIF(TICAM1)-mediated TLR4 signaling
Toll Like Receptor TLR6:TLR2 Cascade
Toll Like Receptor TLR1:TLR2 Cascade
Toll Like Receptor 2 (TLR2) Cascade
MyD88:MAL(TIRAP) cascade initiated on plasma membrane
Toll Like Receptor 3 (TLR3) Cascade
Neutrophil degranulation
Toll Like Receptor 9 (TLR9) Cascade
Axon guidance
Interferon signaling
Signaling by WNT
Cell cycle
Centrosome
Protein signaling
Signaling by GPCR
Disease
epigenetic regulation
lipid metabolism
Interleukin
Cellular responses to external stimuli
Processing of Capped Intron-Containing pre-mRNA
TLR signaling
Astroglia/Choroid
Intermediate Progenitors
Radial glia/ Progenitors
Neuroepithelium
Neurons
Mesenchymal
Retinal ganglion
Miscellaneous
low RNA/ highly infected
Normalized Enrichment Score
2
1
0
-1
-2
Adjusted p-value
0.04
0.03
0.02
0.01


B

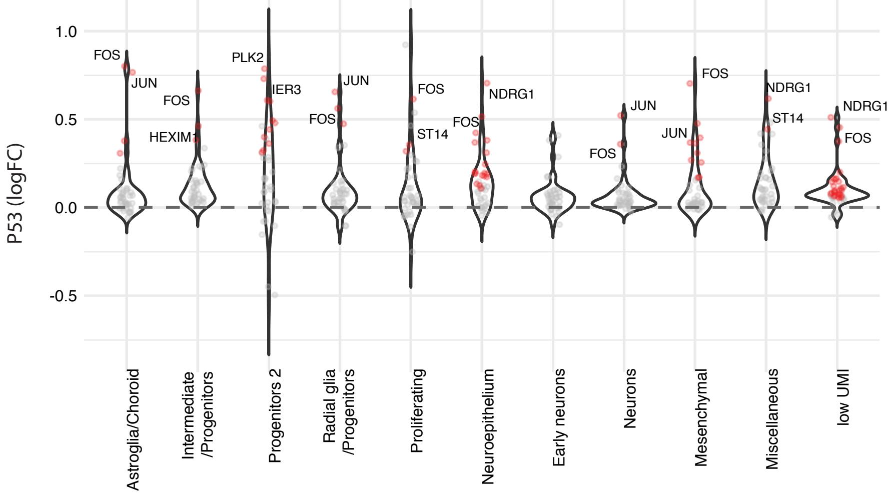

P53 (logFC)
1.0
0.5
0.0
-0.5
FOS
JUN
FOS
HEXIM1
PLK2
IER3
JUN
FOS
FOS
ST14
NDRG1
FOS
JUN
FOS
FOS
JUN
NDRG1
ST14
NDRG1
FOS
Astroglia/Choroid
Intermediate /Progenitors
Progenitors 2
Radial glia /Progenitors
Proliferating
Neuroepithelium
Early neurons
Neurons
Mesenchymal
Miscellaneous
low UMI

**Figure S6.**

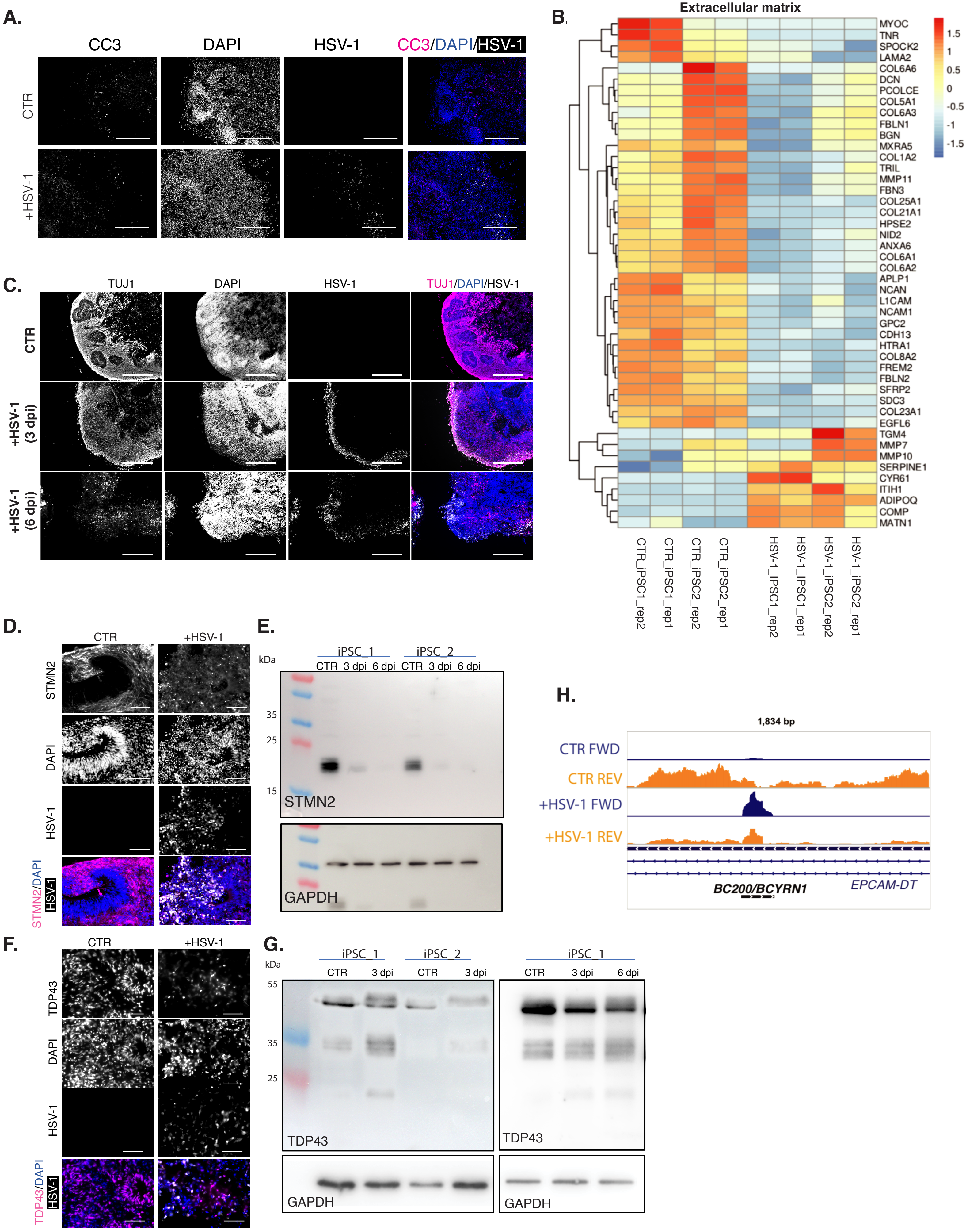
A.
CC3
DAPI
HSV-1
CC3/DAPI/HSV-1
CTR
+HSV-1
B.
Extracellular matrix
MYOC
TNR
SPOCK2
LAMA2
COL6A6
DCN
PCOLCE
COL5A1
COL6A3
FBLN1
BGN
MXRA5
COL1A2
TRIL
MMP11
FBN3
COL25A1
COL21A1
HPSE2
NID2
ANXA6
COL6A1
COL6A2
APLP1
NCAN
L1CAM
NCAM1
GPC2
CDH13
HTRA1
COL8A2
FREM2
FBLN2
SFRP2
SDC3
COL23A1
EGFL6
TGM4
MMP7
MMP10
SERPINE1
CYR61
ITIH1
ADIPOQ
COMP
MATN1
1.5
1
0.5
0
-0.5
-1
-1.5
CTR_iPSC1_rep2
CTR_iPSC1_rep1
CTR_iPSC2_rep2
CTR_iPSC2_rep1
HSV-1_iPSC1_rep2
HSV-1_iPSC1_rep1
HSV-1_iPSC2_rep2
HSV-1_iPSC2_rep1
C.
TUJ1
DAPI
HSV-1
TUJ1/DAPI/HSV-1
CTR
+HSV-1 (3 dpi)
+HSV-1 (6 dpi)
D.
CTR
+HSV-1
STMN2
DAPI
HSV-1
STMN2/DAPI HSV-1
E.
iPSC_1
iPSC_2
kDa
CTR 3 dpi 6 dpi CTR 3 dpi 6 dpi
35
25
15
STMN2
GAPDH
H.
1,834 bp
CTR FWD
CTR REV
+HSV-1 FWD
+HSV-1 REV
BC200/BCYRN1
EPCAM-DT
F.
CTR
+HSV-1
TDP43
DAPI
HSV-1
TDP43/DAPI HSV-1
G.
iPSC_1
iPSC_2
iPSC_1
kDa
CTR 3 dpi CTR 3 dpi
CTR 3 dpi 6 dpi
55
35
25
TDP43
TDP43
GAPDH
GAPDH

#### Figure S7

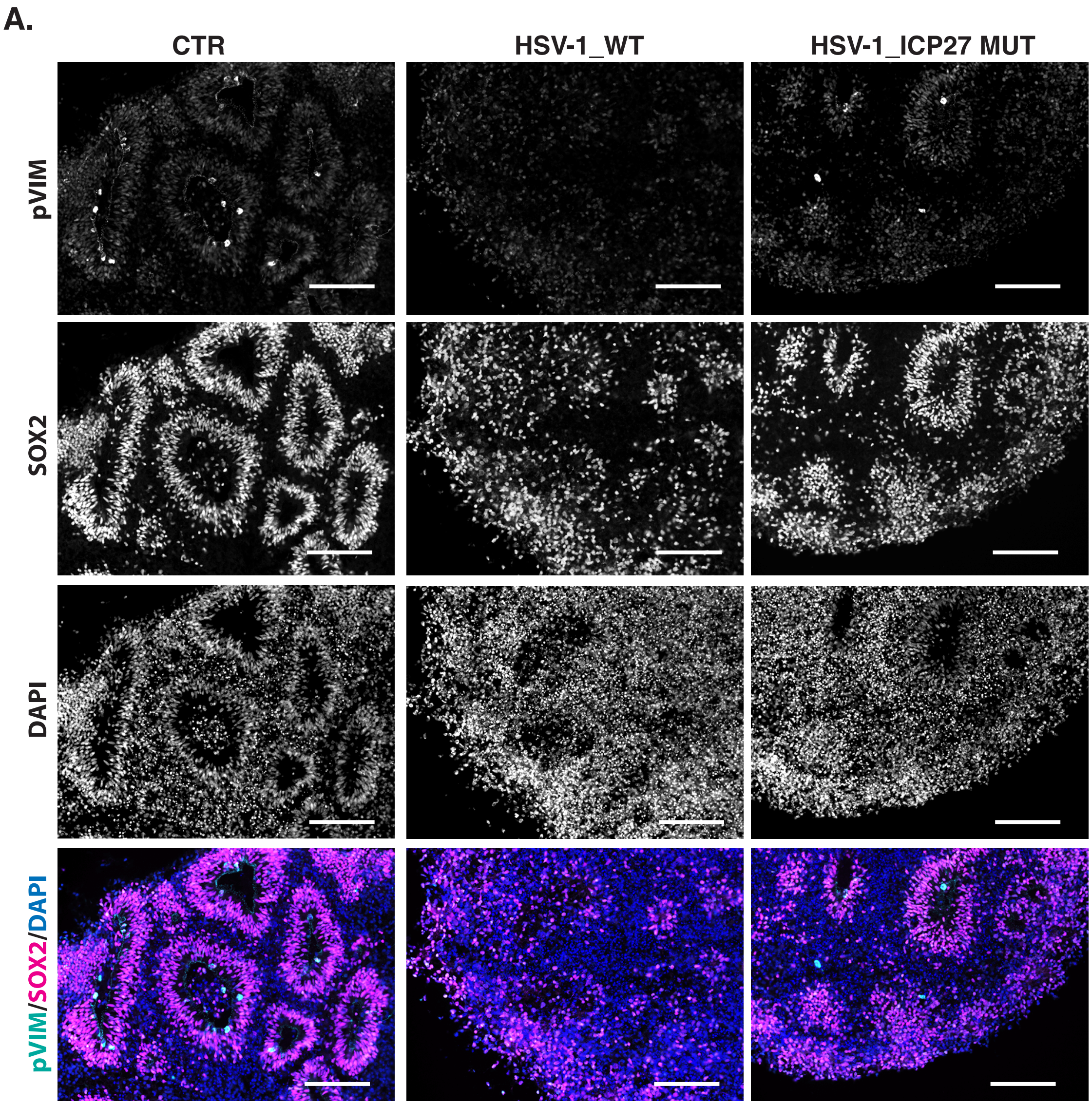

**Supplementary Figures**

**Figure S1. A**. An overview of 30 and 60 days old organoids derived from iPSC line-1 and iPSC line-2 respectively, and immunohistochemistry of organoid cryosections for neuronal progenitors (SOX2, PAX6-red), neurons (TUJ1, MAP2-green), proliferating progenitors (pVIM-green) and astrocytes (GFAP-red). Scale bar - 100 µm. **B.** Quantitative real-time PCR analysis of *TBR1, CTIP2, HOPX and RELN* transcript expression in organoids derived from iPSC line-1, 30, 60 and 90 days of development (three biological replicates). **C.** Nanostring analysis of gene expression in organoids derived from iPSC line-1 and iPSC line-2 at day 0, 30 and 60 of development. **D.** UMAP showing overlap between two biological replicates from control 60 days old brain organoid. **E.** UMAP visualization showing cells colored by their respective cell type in uninfected 30 days old brain organoid. **F.** UMAP plots depicting expression of cell-type specific markers: *DCX*-early neurons, *SPARCL1*-astroglia, MKI67-proliferating radial glia, *SOX2*-progenitors, *EOMES*-intermediate progenitors, *TTR*-choroid plexus. **G.** Representative images from the VISIUM spatial analysis of 60 days old organoids iPSC line-2 for markers: *SYP*-synapses, *SOX2*-progenitors, *DCX*–neurons, *EOMES*-Intermediate progenitors, *MKI67*-proliferating radial glia, *TTR*-choroid plexus. The colors indicate log2 expression. **H.** Schematic overview of experimental set-up. CTR = uninfected, +HSV-1 = infected. **I.** Representative images HSV-1-GFP protein (white) expression in uninfected (CTR) and 3 and 6 days post infection (dpi) 60 days old organoids, DAPI (blue) – stains nuclei. Scale bar - 100 µm.

**Figure S2. A.** A principal component analysis (PCA) of uninfected (CTR) and infected (+HSV-1) organoids derived from iPSC line-1 (red) and iPSC line-2 (blue), (2 biological replicate per sample). **B.** Log2 fold changes of differentially expressed genes from uninfected (CTR) and 6 days infected 60 days old brain organoids from two iPSC lines (n=3 biological replicates per sample). Highlighted genes in green indicate significant changes. **C.** GO term analysis of the top dysregulated transcripts from 6 dpi infected organoids (data combined from two iPSC lines n=2 biological replicates per sample), colors indicate GO terms: yellow – biological process (BP); green – cellular component (CC); red – molecular function (MF). **D.** Heatmap representation of gene expression (GO term chemical synaptic transmission), in two iPSC lines in uninfected (CTR) and 6 days infected (+HSV-1) organoids (2 biological replicates are shown). **E.** Heatmap representation of gene expression (GO term DNA binding/transcription factors), in two iPSC lines in uninfected (CTR) and 6 days infected (+HSV-1) organoids (2 biological replicates are shown). **F.** Plot of the median of firing rate across electrodes for the measured timepoints. Only electrodes with at least 25 spikes at baseline (timepoint -1h) were used to compute the median values. Three independent biological replicates for CTR and 3 for +HSV-1 are shown. Abbreviations: CTR: Control organoids, +HSV-1: Herpes Simplex Virus 1 infected organoids. **G**. Bright field and GFP images of 60 days uninfected (CTR and infected (+HSV-1) organoids placed on multielectrode array. Arrows indicate single cell layer of neuronal processes migrating from organoids. **H.** Line chart showing the progression of the relative activity based on the mean spike frequency of uninfected (CTR) and infected (+HSV-1) after 24 and 48 hours post-infection (hpi). **I.** Mosaic image of the footprint of an uninfected organoid (CTR) and HSV-1-GFP infected organoid (+HSV-1) used in calcium imaging experiments showing TUJ1 expression (red), HSV-1-GFP expression (green) and DAPI (blue). Scale bar – 1mm.

**Figure S3. A.** Scatter plot of log10 normalized read counts along with pairwise Pearson correlations among replicates for cellular (blue) and viral (red) genes **B.** Poly(A) tails length distribution (tail length on x axis, read frequency on y axis) for selected cellular (blue) and all detected viral (red) transcripts. **C**. Cellular poly(A) tails length distribution per mRNA (left panel), per gene (middle panel) in infected (red) and uninfected (blue) organoids. Viral poly(A) tails length distribution per mRNA (right panel). Median lengths are 120,125 and 125 for the three uninfected samples and 125, 141 and 146 for the infected samples. Median viral transcripts tail lengths are 147,178 and 162. Wilcoxon rank sum test between cellular and viral length distributions: p-value = 2.2e-16. Wilcoxon rank sum test between cellular transcripts length distributions in infected vs uninfected: p-value = 2.2e-16. **D.** Box plots of poly(A) tail length per gene (in nucleotides) in uninfected (blue) and infected (red) samples, for genes with significantly longer tails in uninfected (left panel) and significantly longer tails in infected (right panel) samples. **E.** The difference in the change in proximal isoform usage between high readthrough (upper panel) and low readthrough genes (lower panel).

**Figure S4. A.** An overview of cell numbers in each of two replicates of infected (3 and 6 days post infection (dpi)) and uninfected (CTR) organoids. **B.** Violin plots show the log-scaled distribution of gene per cell (left panel); UMIs per cell (middle panel) and percentage of mitochondrial genes (right panel). **C.** UMAP depicting distribution of cells from uninfected (blue) or infected (3 dpi-red and 6 dpi-green) organoids. **D.** UMAP plot depicting expression of cell-type specific markers: *DCX*-early neurons, *HES6-early neurons, MKI67*-proliferating radial glia, *TTR-choroid plexus, STMN2*-neurons, *SPARCL1*-astroglia, *TTR-choroid plexus.* **E**. Representative immunostaining of mitotic marker (pVIM, red) and progenitor marker (SOX2, green) in 60 days old organoids (iPSC line-2) in uninfected (CTR) and infected (+HSV-1) organoids. DAPI (blue) – stains nuclei. Scale bar - 100 µm. **F.** Heatmap showing the expression of the 20 most abundant viral genes (including GFP) in 3 dpi organoid cells. Hierarchical clustering of the cells reveals two distinct 'viral_clusters' based on the expression of early and late viral genes ('viral_cluster' 1) or early genes only ('viral_cluster' 2). Membership to 'final_cluster' and to the 'low_RNA/Highly infected' cluster are displayed as additional annotations, showing that most 'low_RNA/Highly infected' cells are grouped in 'viral_cluster' 2 and show higher expression of multiple early and late viral genes. **G**. Box plot showing normalized expression values for the combined introns of BCL2L11, RASD1, and RRAD per cell. Lower and upper hinges correspond to the first and third quartiles. Whiskers extend to a maximum of 1.5 times the distance between the first and third quartile. Outliers beyond are marked by single dots. Notches extend 1.58 * interquartile range divided by the square root of number of cells. **H.** Representative images of spatial transcriptomics of immediate early-*US1*, early-*UL54* and late-*LAT* gene expression in uninfected (CTR) and 1 and 3 days infected (1 and 3 dpi) 60 days old organoids. The colors indicate log2 expression. **I.** For every cell type, RNA molecule counts (UMI, unique molecular identifiers) from uninfected and 3dpi organoids are displayed boxplots. Median is shown as thick black line, lower and upper hinges correspond to first and third quartiles. Whiskers extend to a maximum of 1.5 times the distance between first and third quartile. Outliers beyond are marked by single dots. **J.** Heatmap of prediction scores for each spatial spot, grouped by capture area, for all 14 clusters detected in the single cell dataset. **K.** Spatial distribution of prediction scores, from 0: no similarity to 1: highest similarity, for the 'Neurons' and 'low-UMI' single cell RNA seq clusters (Figure S5B. and C., respectively) in the spatial transcriptomics datasets. **L.** For each spot in the spatial transcriptomics data from HSV-1 (1 dpi), the GFP expression values are plot on the horizontal axis, and the probabilities for low RNA/highly infected cells (left) or neurons (right) on the vertical axis.

**Figure S5.** **A.** Gene set enrichment analysis (GSEA) on differential expression between uninfected and infected (3 days post infection) organoids for each cell type. Color- Normalized enrichment score (NES) values and dot size -log10 (FDR) from statistically significant pathways (FDR < 0.05). **B.** log2 fold changes of P53 signaling for each cell type. Red (blue) dots: genes with positive (negative) log2 fold changes.

**Figure S6.** **A.** Exemplary western blot analysis of CC3 and housekeeping GAPDH. Right panel: Exemplary immunohistochemistry for CC3 (red) nuclei (DAPI-blue) and HSV-1-GFP (white) in uninfected (CTR) and infected organoids (+HSV-1) in iPSC line-2, scale bar – 100 µm. **B.** Heatmap representation of gene expression (GO term extracellular matrix), in two iPSC lines of uninfected (CTR) and infected (+HSV-1) organoids (2 biological replicates are shown). **C.** Exemplary immunohistochemistry for TUJ1 (neurons, red), HSV-1 (GFP, white) and DAPI (nuclei, blue) in uninfected (CTR) and infected organoids (+HSV-1), 3 and 6 days after infection, scale bar – 100 µm. **D.** Exemplary immunohistochemistry for stathmin-2 (STMN2; neurons, red), TUJ1 (neurons, green), DAPI (nuclei, blue) and HSV-1 (GFP, white) in uninfected (CTR) and infected organoids (+HSV-1) in iPSC line-2, scale bar – 100 µm. **E.** Western blot analysis of STMN2 protein expression in organoids derived from both iPSC lines, GAPDH serves as a loading control. **F.** Exemplary immunohistochemistry for TAR DNA-binding protein 43 (TDP43, red), DAPI (nuclei, blue), HSV-1-GFP (white) in uninfected (CTR) and infected organoids (+HSV-1), iPSC line-2, scale bar – 100 µm. **G.** Western blot analysis of TDP43 protein expression in organoids derived from both iPSC lines, 43 kDa, 35 kDa and 25 kDa isoforms are depicted. GAPDH serves as a loading control. **H.** Coverage profiles from total RNA-sequencing data from 60 days old cerebral organoids for BC200/BCYRN1 transcript in control (CTR) and infected (+HSV-1) organoids. Sense genes in *blue* (orange) oriented *left* to *right (right to left). EPCAM-DT t*ranscript in *dark blue,* BC200/BCYRN1 in black.

**Figure S7. A.** Exemplary immunohistochemistry for pVIM (mitotic marker, green), DAPI (nuclei, blue) and SOX2 (neuronal progenitors, red) in the uninfected (CTR) and organoids infected with wild-type HSV-1 virus (2 days post infection (dpi)) and ICP27 deletion mutant virus (4 dpi), scale bar – 100 µm.

**Video S1. Calcium imaging in 60 days-old organoids.** Time lapse series showing the fluctuations of CalBryte590 fluorescence. Images were acquired at 5 Hz for 5 min and relative changes in intensity over time computed.

**Video S2. Calcium imaging in 60 days-old organoids HSV-1 infected organoid (2 dpi).** Time lapse series showing the fluctuations of CalBryte590 fluorescence. Images were acquired at 5 Hz for 5 min and relative changes in intensity over time computed.
