## Supplementary material for "Neurodegeneration in human brain organoids infected with herpes simplex virus type 1": Table_S3

Primary antibodies for immunofluorescence (IF) and western blotting (WB).

| **Antibodie** | **Isotype** | **Clone** | **IF** | **WB** | **Company** |
|---|---|---|---|---|---|
| CC3 | rabbit | polyclonal | | 1:1000 | Cell Signaling Technology |
| COL3A1 | mouse IgG | polyclonal | | 1:1000 | Novus Biologicals, Littleton, US |
| EMOES | rabbit | polyclonal | | 1:1000 | Abcam, Cambridge, United Kingdom |
| GAPDH | mouse IgM | monoclonal (71.1) | | 1:2000 | Merck-KGaA, Darmstadt, DE |
| GFAP | rabbit | polyclonal | | | Merck-KGaA, Darmstadt, DE |
| ICP0 | mouse | monoclonal | | | Santa Cruz Biotechnology |
| LHX2 | mouse | monoclonal | | 1:100 | DSHB, Iowa, US |
| MAP2 | mouse IgG1 | monoclonal (CL5420) | | 1:1000 | Atlas antibodies, Bromma, SWE |
| PAX6 | mouse | monoclonal | | 1:100 | DSHB, Iowa, US |
| pVIM | mouse, IgG2 | monoclonal (4A4) | | 1:2000 | Medical & Biological Laboratories, Woburn, US |
| RELN | mouse IgG | monoclonal (RE-3B9) | | | DSHB, Iowa, US |
| SOX2 | rabbit IgG | polyclonal | | 1:1000 | Merck-KGaA, Darmstadt, DE |
| SNY1 | rabbit IgG | polyclonal | | 1:1000 | Abcam, Cambridge, United Kingdom, |
| STMN2 | rabbit IgG | polyclonal | 1:1,000 | 1:2000 | Novus Biologicals, Littleton, US |
| TDP-43 | rabbit IgG | polyclonal | 1:1,000 | 1:2000 | Proteintech Group, Manchester, UK |
| TUJ1 | mouse IgG2a | monoclonal (MMS-435P) | | 1:1000 | BioLegend, San Diego, US |

| Primers | Fwd | Rev |
|---|---|---|
| LAT | GATTTCCCGCGTCAATCAGC | GACCCTTACACTGGAACCGG |
| UL29 | GATTTCCCGCGTCAATCAGC | ATGAACAGCTGCAACGGGTA |
| US1 | GATTTCCCGCGTCAATCAGC | ATGAACAGCTGCAACGGGTA |
| COL6A3 | TCTGTTCCTCTTTGACGGCT | GGACTCCACCTTGACATCA |
| BGN | AACAGTGGCTTTGAACCTGG | CAGTTCGATGGCCTGGATTT |
| DES | CCTGGAGCGCAGAATTGAAT | TCATACTGAGCCCGGATGTC |
| HS3ST6 | CTACTTCAACGCCACCAAGG | GTTGAAGGGCCGGTAGAACT |
| HS3ST2 | CCCCACTTCTTTGACAGGAA | TGTCTCGGGACATGTTGAAG |
| TBR1 | ACTGGCTCGAGCGACTTTTA | CCGATTTCCTTACTCCACCA |
| RELN | CAGGCAACTGGCTTTTCTTC | CATCCTGTAGGCTGGCTCTC |
| GAPDH | AAGGTGAAGGTCGGAGTCAAC | GGGGTCATTGATGGCAACAATA |
| GFAP | CCACCTACAGGAAGCTGCTA | TGGCCTTCTGACACAGACTTG |
| SYP | ATTGTGCCAACAAGACCGAGAGT | CAGGAAGATGTAGGTGGCCAGAG |
